# Biophysical forces rewire cell metabolism to guide microtubule-dependent cell mechanics

**DOI:** 10.1101/2020.03.10.985036

**Authors:** Stephanie Torrino, Stephane Audebert, Ilyes Belhadj, Caroline Lacoux, Sabrina Pisano, Sophie Abélanet, Frederic Brau, Stephen Y. Chan, Bernard Mari, William M Oldham, Thomas Bertero

## Abstract

Mechanical signals regulate cell shape and influence cell metabolism and behavior. Cells withstand external forces by adjusting the stiffness of its cytoskeleton. Microtubules (MTs) act as compression-bearing elements in response to mechanical cues. Therefore, MT dynamics affect cell mechanics. Yet, how mechanical loads control MT dynamics to adjust cell mechanics to its locally constrained environment has remained unclear. Here, we show that mechanical forces rewire glutamine metabolism to promote MT glutamylation and force cell mechanics, thereby modulating mechanodependent cell functions. Pharmacologic inhibition of glutamine metabolism decreased MT glutamylation and affected their mechanical stabilization. Similarly, depletion of the tubulin glutamylase TTLL4 or overexpression of tubulin mutants lacking glutamylation site(s) increased MT dynamics, cell compliance and contractility, and thereby impacted cell spreading, proliferation and migration. Together our results indicate that mechanical cues sustain cell mechanics through glutaminolysis-dependent MT glutamylation, linking cell metabolism to MT dynamics and cell mechanics. Furthermore, our results decipher part of the enigmatic tubulin code that coordinates the fine tunable properties of MT mechanics, allowing cells to adjust the stiffness of their cytoskeleton to the mechanical loads of their environment.

## Main

Fundamental aspects of cell behavior in living organisms—morphogenesis, collective migration and self-organization—are emergent properties of cells interconnected within a tissue network. Understanding the spatiotemporal control of cell behavior thus requires incorporation of information on how structural and architectural complexity of tissues is transmitted to their constituent cells^1^. Mechanotransduction enables cells to sense and adapt to external forces^2^. This mechanical response of cells involves the rapid remodeling of their cytoskeleton and regulates their metabolic states, which are dynamically transited to match energetic and biosynthetic requirements^3–6^. While, major emphasis in the mechanotransduction field has been placed on actin, the dynamics of microtubules (MTs) in response to mechanical cues suggests an equally important role for tubulin^7^.

MTs are long, stiff polymers of αβ-tubulin that are structurally and functionally important components of the eukaryotic cell cytoskeleton, forming the mitotic spindle and the axonemes of cilia and flagella and serving as tracks for intracellular trafficking^8^. Yet, implications of MTs in the mechanotransduction processes are only emerging, and how MT dynamics affect mechanotransduction remains elusive.

To determine whether mechanical cues conveyed by extracellular matrix (ECM) stiffness modulate MT dynamics, we performed fluorescence recovery after photobleaching experiments (FRAP, **Figure 1a** and **Extended Figure 1a**) and monitored growing microtubule plus ends (EB1-EGFP time laps imaged, **Figure 1b**) on HeLa cells cultivated on a gradient of matrix stiffness. ECM stiffening increased the diffusion rate and decreased the mobile fraction of tubulin (**Figure 1a** an **Extended Figure 1a**), while decreased MT growth rate (**Figure 1b**). Together, these results indicate that ECM stiffening stabilize MT. Consistent with previously published results^7,9^, decreasing MT dynamics via taxol decreased cell contractility, while MT destabilization by nocodazole increased cell contractility, as measured by traction force microscopy **(Extended figure 1b**). Thus, ECM mechanical cues influence cell mechanics through MT dynamics.

**Figure 1:**
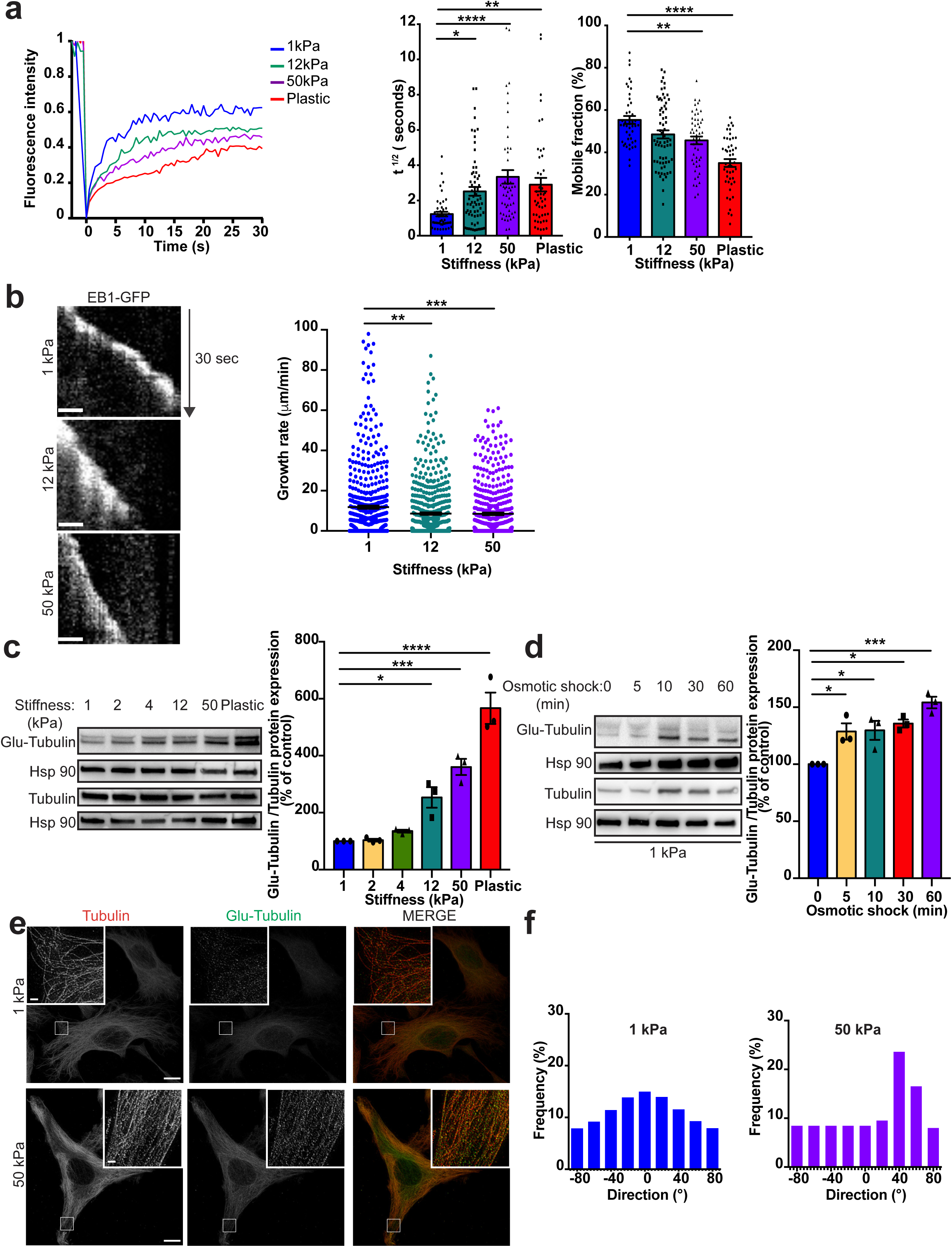
Mechanical cues force microtubules glutamylation to stabilize the microtubule lattice. **(a)** Representative FRAP curves (left) and quantification of diffusion rate (t ½) and mobile fraction (right) of GFP-Tubulin (n>45 cells) **(b)** Representative kymographs (left) and growth rates quantification (right) of EB1-GFP in cells plated on the indicated substrate. n>500 comets. Scale bar=1 µm. **(c-d)** Immunoblot and quantification of Glu-Tubulin in cells plated on the indicated substrate **(c)** or after hypo-osmotic shock (**d). (e-f)** Representative STED images of Tubulin and Glu-Tubulin localization **(e)** and representative alignment of microtubule **(f)** in cells plated on 1 or 50 kPa hydrogel (n>10). Scale bar=10 µm; for the inset, scale bar=1 µm.; *P<0.05; **P<0.01; ***P<0.001; ****P<0.0001; **(a-d)** Bonferroni’s multiple comparison test; data are mean ± s.e.m of at least n=3 independent experiments.

MTs posttranslational modifications (PTMs) have been previously shown to modulate MT persistence^10–12^. Tubulin glutamylation has been reported to control MT severing^13^. Thus, we investigated whether mechanical forces modulate MT glutamylation (**Figure 1c-d** and **Extended figure 1c-h**). As quantified by immunoblotting and immunofluorescence, ECM stiffening (**Figure 1c** and **Extended figure 1f**), osmotic shock (**Figure 1d** and **Extended figure 1h**), and circular shear stress (**Extended figure 1c**) independently promoted MT glutamylation. Importantly, mechano-induced MT glutamylation was also observed in a breast cancer cell line and primary cells (**Extended figure 1d-e**). Furthermore, and consistent with previously published results^14–17^, upon mechanical stresses the MT lattice was robustly reorganized (**Figure 1e-f and Extended figure 1f-h**). More specifically, mechanical forces switched MT organization from a net-like phenotype to alignment via cortical arrays. Thus, mechanical cues increase MT glutamylation and reorganize the MT lattice.

Upon mechanical stress, the large increase of MT glutamylation is likely to require the mobilization of an important intracellular pool of glutamate, especially to ensure the persistence of the phenotype. Recently, we reported that mechanical cues promote glutamine catabolism, a process --mediated by the glutaminase (GLS) to sustain the metabolic needs of mechanoactivated cells^3,5^. Thus, we hypothesized that mechanical loads increase glutamine uptake and catabolism to fuel the intracellular pool of glutamate required for MT glutamylation. Consistent with our hypothesis, steady state metabolomics profiles of HeLa cells indicated that both ECM stiffening (**Extended Figure 2a**) and osmotic shock (**Extended Figure 2b**) promote substantial global alterations of metabolism (**Extended Figure 2a-d**). Steady state levels of 33 metabolites highly enriched in pathways related to amino acid metabolism (aspartate, glutamate, arginine and proline) were significantly changed by both ECM stiffening and osmotic shock (**Figure 2a-b**). These analyses revealed a significant increase in the intracellular glutamate/glutamine ratio (**Extended Figure 2e-f**), while HeLa cells increased their rate of glutamine uptake upon matrix stiffening (**Extended figure 2g**). Importantly, increased GLS expression (**Extended figure 2h)** and activity (**Extended figure 2i**) was observed in cells cultivated on stiff matrix. Taken together, these results support the notion that glutamine uptake and catabolism regulates the level of intracellular pool of glutamate in HeLa cells to sustain MT glutamylation under mechanical stresses.

**Figure 2:**
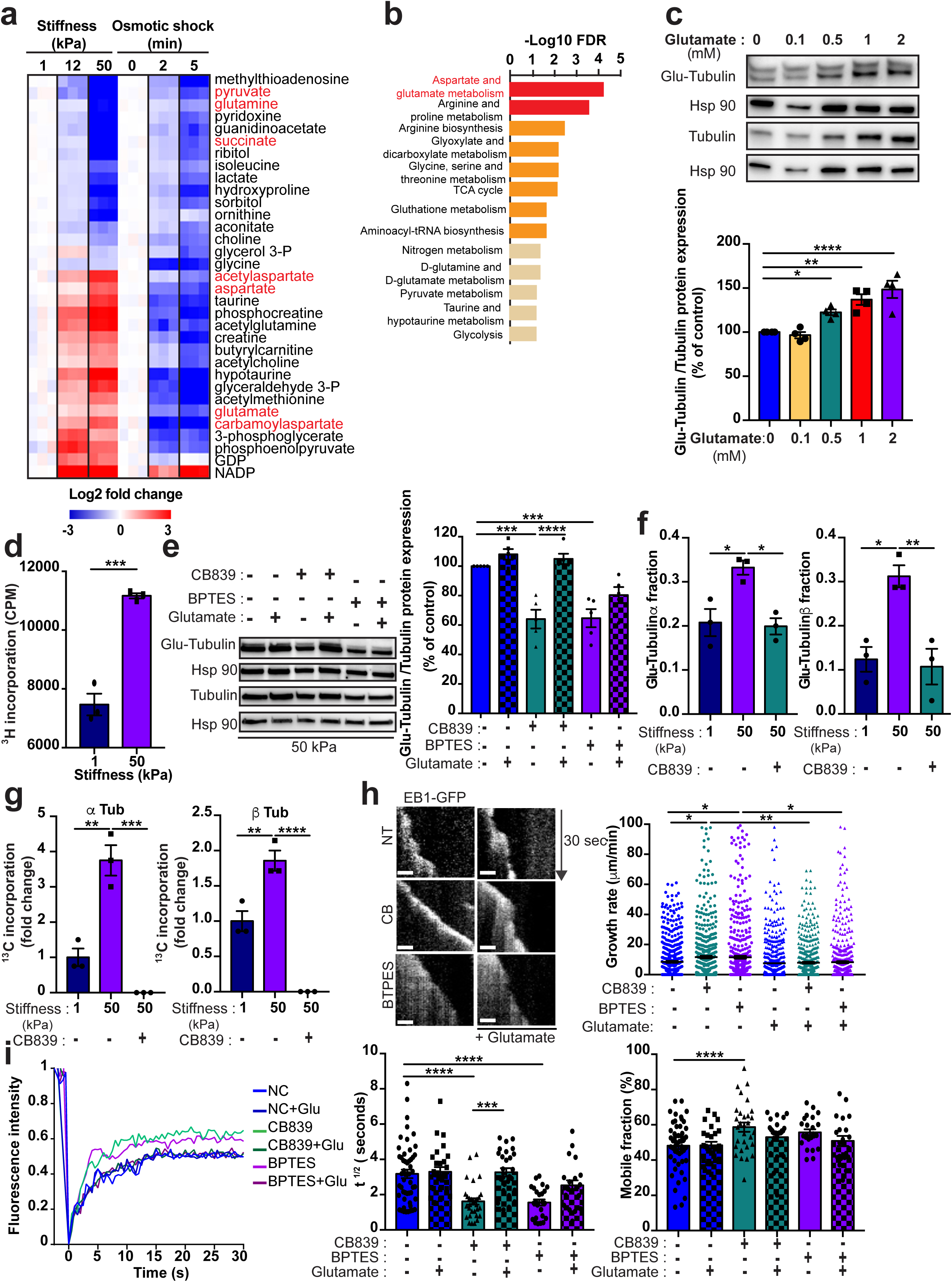
Microtubule glutamylation rely on glutamine catabolism. **(a-b)** Heatmap **(a)** and pathway enrichment analysis **(b)** of significantly (FDR<1%; P<0.05) modulated intracellular metabolites in cells followed indicated treatments. Red: metabolite from aspartate glutamate pathway **(c)** Immunoblot and quantification of Glu-Tubulin in SLC1A3 overexpressing cells plated on 1kPa hydrogel in presence of glutamate. **(d)** ^3^H-glutamine incorporation in purified microtubule from cells treated as indicated in presence of cycloheximide. **(e-i)** Cells plated on the indicated substrate and treated with CB839 or BPTES. **(e)** Immunoblot and quantification of Glu-Tubulin in cells in presence of glutamate. **(f)** Proteomic quantification of glutamylated C-terminal α- and β-tubulin tails in purified microtubule from cells treated as indicated. **(g)** Quantification of ^13^C-glutamine incorporation by stable carbon isotope proteomic in purified microtubule from cells treated as indicated in presence of cycloheximide. **(h)** Representative kymographs (left) and growth rates quantification (right) of EB1-GFP in presence of glutamate n>500 comets. Scale bar=1 µm. **(i)** Representative FRAP curves (left) and quantification of diffusion rate (t ½) and mobile fraction (right) of GFP-Tubulin (n>20) in presence of glutamate.*P<0.05;**P<0.01; ***P<0.001; ****P<0.0001; **(c**,**e-g**,**i)** Bonferroni’s multiple comparison test; **(d)** two tailed t-test; data are mean ± s.e.m of at least n=3 independent experiments.

To investigate whether the level of intracellular pool of glutamate is key to promote MT glutamylation, SLC1A3 overexpressing cells were cultivated on soft substrate in DMEM high glucose, 10% serum, 2mM glutamine and exposed for five minutes to various concentration of glutamate. Increased extracellular glutamate concentration increased intracellular glutamate (**Extended figure 2j**) and MT glutamylation (**Figure 2c**) indicating that, intracellular glutamate levels control MT glutamylation.

We next cultivated HeLa cells on soft vs. stiff substrate and exposed to 200mCi/mL of ^3^H-glutamine for 2 hours in presence of cycloheximide -- a translational elongation inhibitor. As revealed by scintigraphy, increased ^3^H incorporation in purified MT was observed in cells plated on stiff substrate, suggesting that glutamine support MT glutamylation. Then we investigated whether either pharmacological inhibition of GLS -- BPTES [bis-2-(5-phenylacetamido-1,3,4-thiadiazol-2-yl)ethyl sulphide] or CB839, or genetic inhibition of GLS (siGLS) impacted MT glutamylation.Inhibition of GLS using the two independent approaches blunted stiffness-induced glutamine catabolism, as quantified by LC-MS (**Extended figure 3**). and strongly decreased MTs glutamylation, as quantified by immunoblot (**Figure 2e** and **Extended Figure 4a-c)** and proteomic analyses of purified MT (**Figure 2f**). Strikingly, upon stiff matrix (50kPa), the GLS inhibitor-dependent decrease of MT glutamylation was associated with reorganization of the MTs networks to a net-like phenotype (**Extended Figure 4d-e**), phenocopying the observed events in cells cultivated on soft matrix (1kPa; **Figure 1d-e**). Importantly, both MT glutamylation (**Figure 2e** and **Extended Figure 4c**) and MT alignment (**Extended Figure 4d-e**) were rescued by glutamate supplementation.

To definitively establish the critical role of glutamine-GLS-glutamate axis for MT glutamylation, we performed ^13^C_5_-glutamine tracing experiments coupled with proteomic analyses of purified MTs (**Figure 2g**). Cells were cultivated on soft or stiff substrate and exposed to 2mM ^13^C_5_-glutamine for two hours in presence of cycloheximide. Increased ^13^C-glutamate incorporation on both α-tubulin and β-tubulin C-terminal tails was observed in cells cultivated on stiff substrate. Importantly, CB839 treatment abolished MT new glutamylation.

We next investigated whether GLS-dependent MT glutamylation modulates MT dynamics. On stiff matrix (50kPa), GLS inhibition increased MT growth rate (**Figure 2h**) and decreased tubulin diffusion rate while increasing the mobile fraction (**Figure 2i**). Importantly, glutamate supplementation rescued MT stability (**Figure 2h-i**). Thus, decreasing MT glutamylation in cells cultivated on stiff substrate decreased their stability. Taken together, these results demonstrate that mechanical stresses rewire cell glutamine metabolism to sustain the glutamate production required for MT glutamylation-dependent stabilization.

To gain insight into the molecular mechanisms associated with this process, we next sought to identify the enzymes involved in tubulin glutamylation following mechanical stress. MT glutamylation is a dynamic process that occurs through a balance between glutamate addition by a family of glutamylase enzymes (TTLLs)^18^, and glutamate removal by a family of deglutamylase enzymes (CCPs)^19,20^. We performed siRNA screening to identify TTLL and CCP enzyme family members responsible for MT glutamylation in response to mechanical stress (**Figure 3** and **Extended figure 5**). Knockdown efficiency was first validated by RT-qPCR (**Extended figure 5a**). We found that TTLL4, and, to a lesser extent, TTLL5 and TTLL9 depletion significantly decreased MT glutamylation (**Figure 3a**), while CCP5 was the sole deglutamylase where knockdown robustly increased MT glutamylation (**Extended Figure 5b**). These results were confirmed using three distinct single siRNAs directed against TTLL4 and CCP5 (**Extended figure 5c-f**).

**Figure 3:**
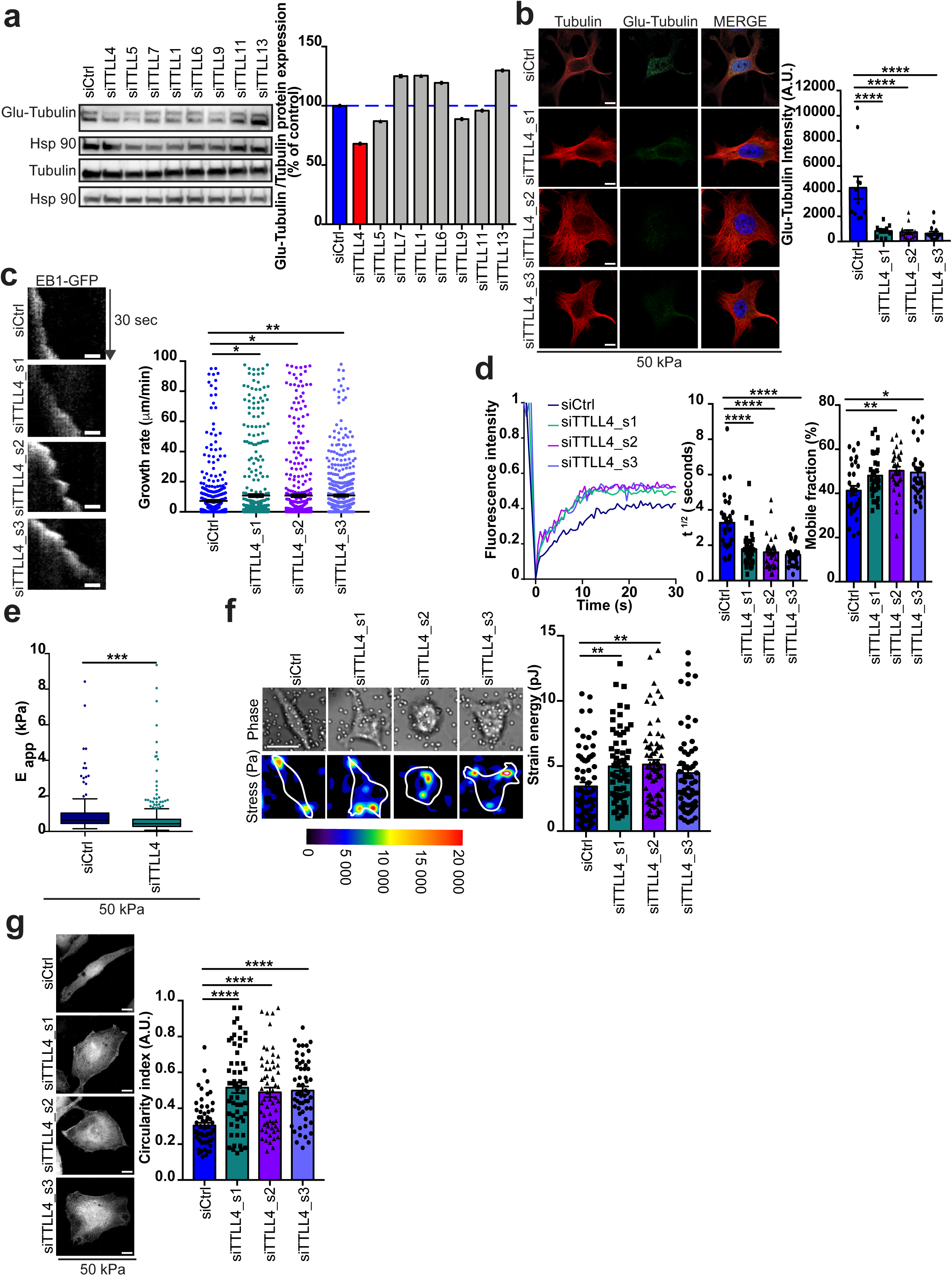
TTLL4 force microtubule glutamylation to adjust cell mechanics and sustain cell mechanic-dependent activities. **(a-f**,**h)** HeLa cells plated on 50 kPa hydrogel were transfected with the indicated siRNA. **(a)** Immunoblot and quantification of Glu-Tubulin in cells. **(b)** Representative confocal images (left) and quantification (right) of Glu-Tubulin and Tubulin. Scale bar=10 µm. **(c)** Representative kymographs (left) and growth rates quantification (right) of EB1-GFP. n>500 comets. Scale bar=1 µm. **(d)** Representative FRAP curves (left) and quantification of diffusion rate (t ½) and mobile fraction (right) of GFP-Tubulin. **(e)** Apparent Young’s moduli obtained by AFM analysis. Bars represent the median. **(f)** Representative heat map (left) and quantification (right) showing contractile forces generate by cells. **(g)** Representative confocal images (left) and quantification (right) of circularity index. Scale bar=10 µm. In all the panels n>20 cells from 3 independent experiments were analyzed. *P<0.05; **P<0.01; ***P<0.001; ****P<0.0001; **(b-d**,**g)** Bonferroni’s multiple comparison test; **(f)** two tailed t-test; data are mean ± s.e.m.

We next investigated whether modulating these enzymes affected the MT lattice. Consistent with our immunoblot results, confocal microscopy confirmed that siRNA knockdown of TTLL4 decreased tubulin glutamylation (**Figure 3b**). Strikingly, decreased MT glutamylation in cell cultivated on stiff matrix (50kPa) reorganized the MT networks to a net-like phenotype (**Figure 3b** and **Extended figure 5g**), as observed in cell cultivated on soft matrix (1kPa). Conversely siRNA knockdown of CCP5 in cells cultivated on soft matrix (1kPa) increased MT glutamylation (**Extended Figure 5h**) and aligned MTs (**Extended figure 5i**) as observed in cell cultivated on stiff matrix. Then, we monitored growing microtubule plus ends and performed FRAP experiments, thus finding that modulation of MT glutamylation impacted MT dynamics in cells (**Figure 3c-d and Extended figure 6a-b**). Altogether, we establish that in response to matrix stiffening, MT glutamylation is orchestrated by TTLL4 and CCP5 which ensure MT organization. Forced MT glutamylation in cells cultivated on soft matrix is sufficient to induce a stiff-like MT lattice phenotype, while MT glutamylation on stiff matrix is necessary to sustain the phenotype. Thus, MT glutamylation is necessary and sufficient to organize the MT lattice.

We next interrogated the importance of MT glutamylation and subsequent MT lattice organization on cell mechanical properties and associated-cellular functions (**Figure 3e-h, Extended figure 6** and **Extended Figure 7**). To investigate whether MT glutamylation impacts cell elasticity, cell contractility and cell membrane tension, cells were plated on either soft (1kPa; siCCP5) or stiff matrix (50kPa; siTTLL4), and we performed atomic force microscopy (**Figure 3e**) and traction force microscopy experiments (**Figure 3f** and **Extended figure 6c**). We observed increased cell compliance (**Figure 3e**) and increased cell traction (**Figure 3f**) in TTLL4-depleted cells, while converse observations were detected in CCP5-depleted cells (**Extended figure 6c**), demonstrating that MT (de)glutamylation affects cell mechanical properties.

On the basis of these findings, we investigated whether tubulin glutamylation may also impact cell functions dependent upon biophysical mechanics such as cellular shape, adhesion, proliferation and migration. Using a similar experimental design, we found that forced tubulin deglutamylation by knocking down TTLL4 (siTTLL4), TTLL5 (siTTLL5) or TTLL9 (siTTLL9) on stiff matrix (50kPa) increased the cell circularity index (**Figure 3g** and **Extended Figure 7**) while decreased both cell proliferation (**Extended Figure 6e and Extended Figure 7**) and migration (**Extended Figure 6i and Extended Figure 7**). Conversely, forced tubulin glutamylation on soft substrate (1kPa) by knocking down CCP5 (siCCP5) decreased the cell circularity index (**Extended Figure 6f**), increased cell proliferation (**Extended Figure 6h**), and decreased cell migration (Extended **Figure 6i**). Modulation of MT glutamylation had no effect on cell adhesion (**Extended figure 6g-h)**. Together, our results indicate that (de)glutamylation enzymes play a central role on cell mechanical properties and dependent cell functions.

To demonstrate the central role of tubulin (de)glutamylation on MT dynamics, cell mechanics, and mechanodependent cell functions, we investigated MT lattice organization, MT dynamics, and cell mechanics of HeLa cells overexpressing tubulin mutants lacking glutamylation site(s) (**Figure 4** and **Extended Figure 8**). As quantified by immunoblotting (**Figure 4b**) and immunofluorescence (**Figure 4c**) forced expression of either Tub ΔCter (a tubulin lacking the eleven C-terminal amino acid) or Tub_E445D (a tubulin in which the glutamate at position 445 is mutated to aspartate^26–29^) decreased MT glutamylation. Consistent with our previous results, decreased MT glutamylation reorganized the MT lattice (**Figure 4c** and **Extended Figure 8a**), modulated MT dynamics (**Figure 4d** and **Extended Figure 8b**), increased cell compliance and cell contractility (**Figure 4e-f**), increased cell circularity (**Figure 4g**) and decreased both cell proliferation and migration (**Extended Figure 8c-d**). Together, our results underline the central role of tubulin (de)glutamylation on cell mechanical properties and cell behavior, including migration and proliferation.

**Figure 4:**
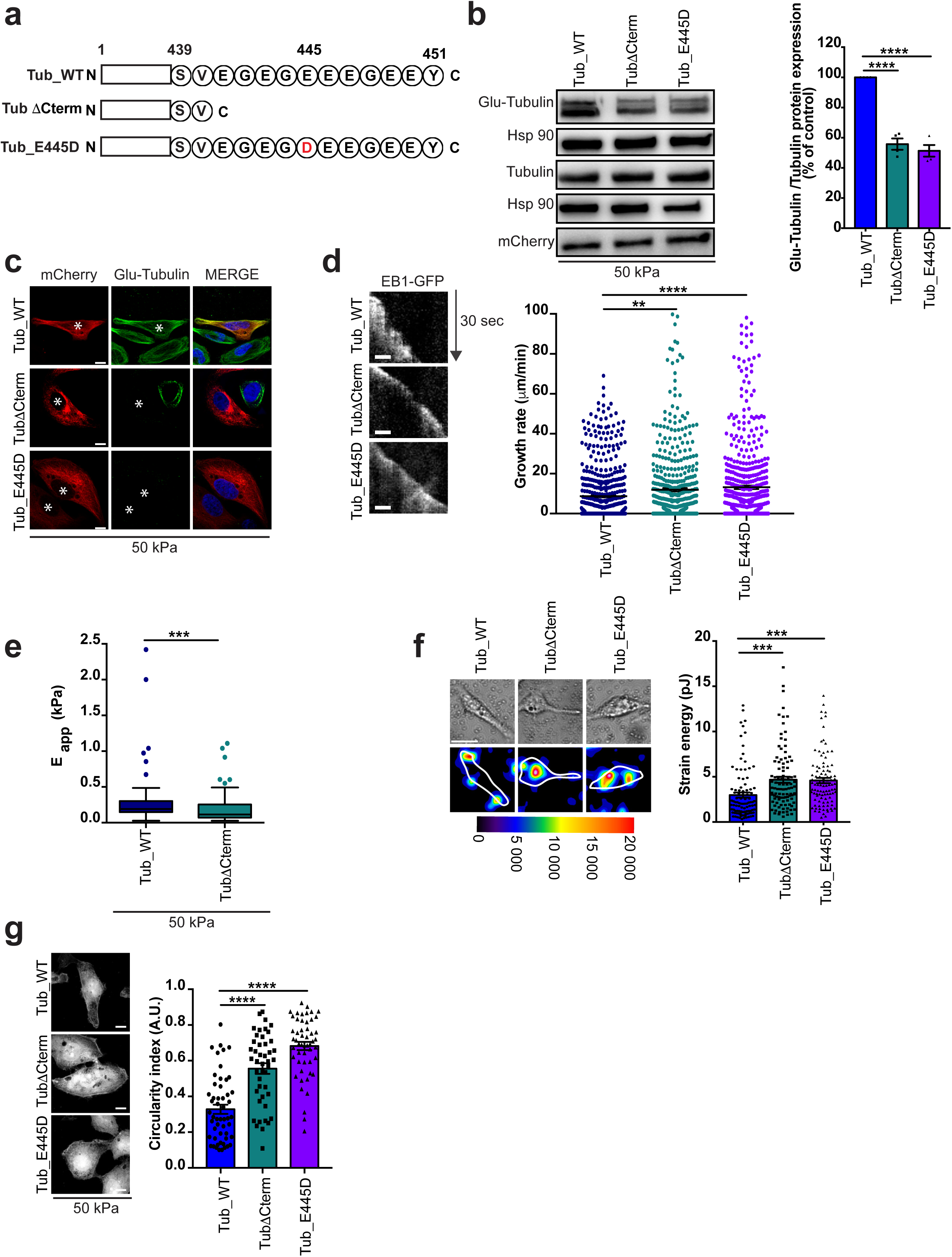
Microtubule glutamylation is sufficient to adjust cell mechanics and sustain cell mechanic-dependent activities. **(a)** Schematic representation of TUBA1A structure in wild type and mutant. (b-g) HeLa cells were transfected with TUBA1A constructs. **(b)** Immunoblot and quantification of Glu-Tubulin in cells. **(c)** Representative confocal images (left) and quantification (right) of Glu-Tubulin and Tubulin. Scale bar=10 µm. *: Transfected cell **(d)** Representative kymographs (left) and growth rates quantification (right) of EB1-GFP. n>500 comets. Scale bar=1 µm. **(e)** Apparent Young’s moduli obtained by AFM analysis. Bars represent the median. **(f)** Representative heat map (left) and quantification (right) showing contractile forces generate by cells. **(g)** Representative confocal images (left) and quantification (right) of circularity index. Scale bar=10 µm. In all the panels n>50 cells from 3 independent experiments were analyzed. **P<0.01; ***P<0.001; ****P<0.0001; **(b, f-g)** Bonferroni’s multiple comparison test; **(e)** two tailed t-test; data are mean ± s.e.m.

Overall, we show here that the mechano-dependent metabolic rewiring of cells represents a critical mediator of MT mechanics. While previous studies demonstrated the role of MT dynamics in cell mechanics^7^, the potential significance of MT PTMs and their associated dependence on cell metabolism has not been described previously. We demonstrate that mechanical cues precisely coordinate cell metabolism and MT glutamylation to stabilize the MT lattice in order to adjust cell cytoskeleton stiffness and adapt to mechanical load. Although several mechanotransduction pathways have been uncovered^2,30–33^, the molecular mechanisms identified so far have mainly addressed the perception of stress intensity and not the direction of the mechanical information. Here, by uncovering MTs glutamylation as a necessary and sufficient PTM to reorganize the MTs lattice under mechanical stresses and to increase MTs persistence, we point out MTs as ideal anisotropic sensors to perceive the directions of cell-scale mechanical signals^34,35^. Yet, whether other partners are needed for MTs to sense tension remain to be addressed, and further studies should evaluate the importance of cytoskeletal crosstalk^22^ in orchestrating cell mechanics in a precise and highly adaptable manner.

Our study also uncovered the PTM enzymes involved in this specific pathway. Previous studies linking MT dynamics to cell mechanics mostly relied on the use of MT-targeting agents^7,9,36^ such as taxol and nocodazole that markedly disrupt MT dynamics and profoundly disorganized the MT lattice. Here, we identify the specific enzymes involved in tubulin glutamylation in response to mechanical cues. By identifying TTLL4 and CCP5 as crucial mediators of this process, we unveil the fine tunable properties of MT mechanics allowing cells to adjust the stiffness of their cytoskeleton to the mechanical loads of their environment. Because TTLL4 has recently been shown to glutamylate several other substrates^37–39^, we investigated the cell mechanical properties of cells expressing tubulin mutants, demonstrating that TTLL4-regulated cell mechanics strictly rely on MT glutamylation. Yet, the true breadth of influence of MT PTMs on cell mechanics and mechanodependent cellular function may involve additional molecular components. While here we reported that MT glutamylation stabilize MT, our results showing that siRNA knockdown of TTLL13 increased MT glutamylation (**Figure 3a**) and siRNA knockdown of CCP4 decreased MT glutamylation (**Extended Figure 5b**) suggest a previously reported biphasic dependence of MT stability on glutamylation^13^. Further studies are necessary to investigate the physiological relevance of this mechano-induced feedback loop. In addition, while we demonstrated the importance of MT glutamylation on several cell mechanodependent cell functions such as proliferation and migration in 2 dimensional cell culture, further studies are necessary to decipher the role of MT glutamylation in 3 dimensional space, thereby refining the molecular codes embedded in this complex network of post-translationally modified tubulin isoforms.

## Methods

### Reagents and antibodies

Nocodazole (487928), Taxol (T7402), Cycloheximide (01810), BPTES (SML0601) were purchased from Sigma-Aldric and were used at the final concentration of 10µM. Glutamate (5mM) was purchased from Gibco. CB839 (S76655) was purchased from Selleck Chemicals and used at the final concentration of 1µM. MES buffer (15424169), GTP (1mM; 15255116), DTT (10699530), Protease and Phosphatase Inhibitor (15662249) were purchased from Thermo Fisher Scientific. The following commercially available antibodies were used for western blotting and immunofluorescence: - mouse monoclonal antibodies against Glu-Tubulin (CliniSciences, AG-20B-0020-C100), Tubulin (Santa Cruz Biotechnology, sc-398937), Hsp90 (Santa Cruz Biotechnology, sc-69703), Hsp60 (Santa Cruz Biotechnology, sc-271215); - rabbit polyclonal antibodies against Tubulin (Sigma-Aldrich, SAB4500087), TTLL4 (Bio-Techne, NBP1-81535), CCP5 (Abcam, ab170541), GLS (Abcam, ab93434). HRP-conjugated donkey anti-mouse IgG (715-035-150) and HRP-conjugated anti-mouse IgG (711-035-152) were purchased from Jackson ImmunoResearch Laboratories.

### Cell culture

HeLa cells and MDA-MB-468 cells were purchased from the American Type Culture Collection (ATCC). Cells used in this study were within 20 passages after thawing and were cultured (37°C, 5% CO_2_) in Dulbecco’s Modified Eagle’s Medium (DMEM, Gibco) supplemented with 10% fetal bovine serum (Gibco), Glutamine (2mM, Gibco) and Penicillin/Streptomycin (1%, Gibco). Primary human pulmonary arterial smooth muscle cells (PASMCs) were cultured in SmGM-2 cell culture media (Lonza), and experiments were performed at passages 3 to 9. For the studies dependent on matrix stiffness, collagen-coated hydrogel pre-plated in culture wells (Matrigen) was generated from a mix of acrylamide and bis-acrylamide coated with collagen. Cells were cultured, passaged, and harvested while on top of the hydrogel, using standard cell culture techniques.

### siRNA and plasmid transfection

Cells were plated on collagen-coated plastic (50µg/mL) and transfected 24h later at 70-80% confluence using siRNA (25nM) and Lipofectamine 2000 reagent (Life Technologies), according to the manufacturers’ instructions. Eight hours after transfection, cells were trypsinized and re-plated on hydrogel. The PMXS retroviral vector containing the coding sequence for SLC1A3 was purchased (Addgene; Plasmid #72873^40^). The mammalian expression vector containing the coding sequence for EB1-2xEGFP was purchased (Addgene; Plasmid # 37827^41^) The pShuttle mCherry-Tubulin adenoviral vector (Tub-WT) was purchased (Addgene; Plasmid # 26768^42^). Tubulin mutants have been generated using the pShuttle mCherry-Tubulin plasmid following recommendations of the Q5 site directed-mutagenesis kit from New England Biolab (#E0554). Oligos Forward pShut: GCGCCGCTCGAGCCTAAG and Reverse TubΔCter: TTAAACAGAATCCACACCAACCTCCT were used for the Tubulin deletion mutant (TubΔCter) and oligos Forward pShut: GCGCCGCTCGAGCCTAAG and Reverse TubE445D: TTAGTATTCCTCTCCTTC**A**TCCTCACCCT were used for the Tubulin Substitution mutant (TUBE445D). pShuttle mCherry-Tubulin mutations were verified by sequencing using oligos pASV40: GAAATTTGTGATGCTATTGC and SeqTub: CAGGTCTCCACCAGGCACCA.

Forty-eight after transfection cells were harvest for analysis. siRNA ON-TARGETplus Human or Non-Targeting Control siRNAs (D-001810-01) were purchased from Horizon Discovery Ltd). Sequences of SMARTpool or single siRNA ON-TARGETplus Human are provided in the Table:

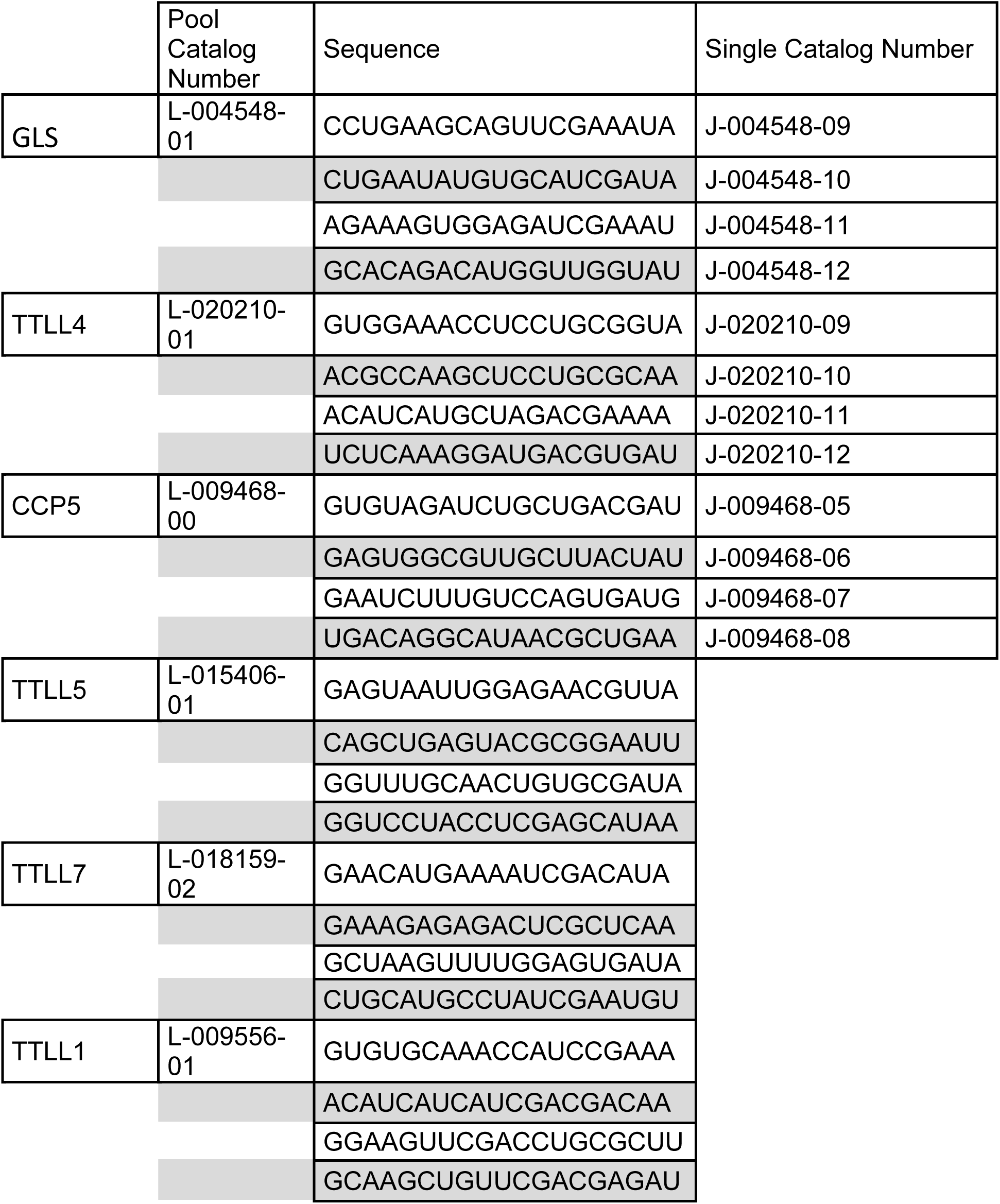
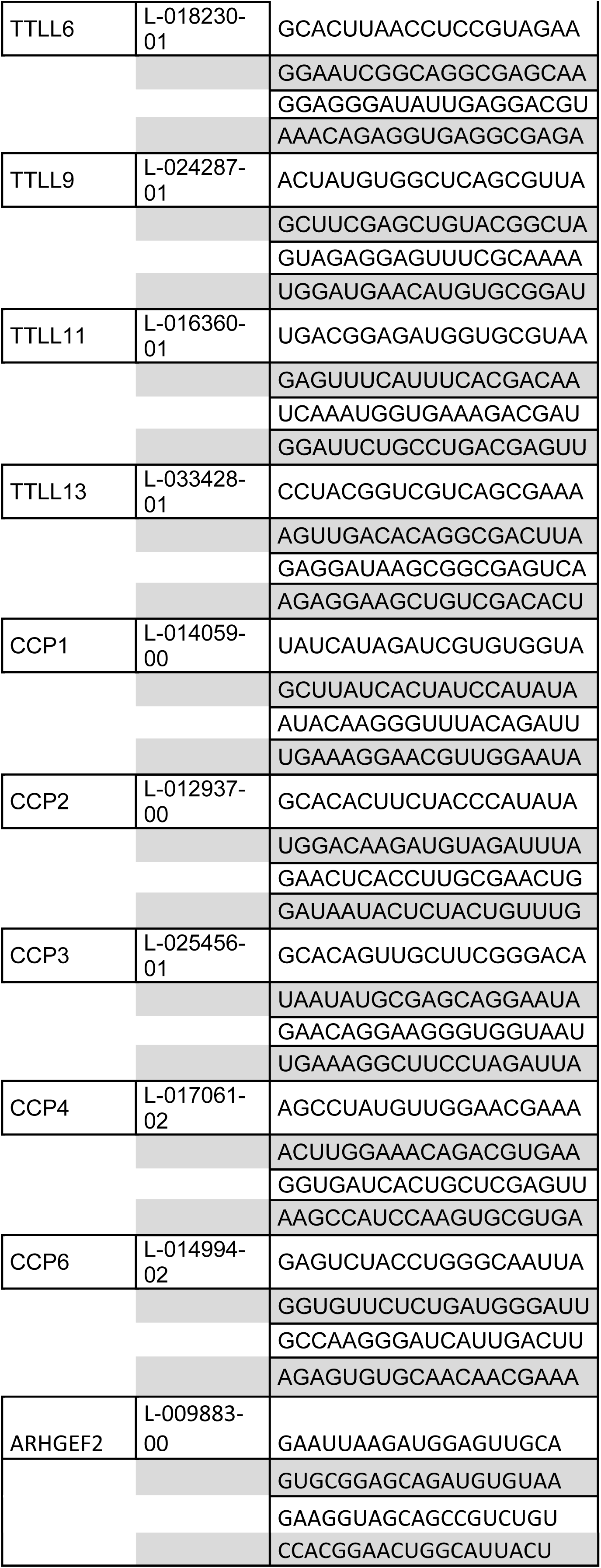

### Immunofluorescence

Cells were fixed with PBS/PFA 4% for 10 min and permeabilized with PBS/Triton 100X 0.2% for 5 min. After blocking with PBS/Triton/BSA 0.2% for 1h, the cells were then incubated with primary antibodies (1/100) at room temperature for 1 h. Secondary antibodies coupled with Alexa-594 and/or Alexa-488 (A-11012, A-11001, Life technologies) were used at 1/500 for confocal imaging, while Alexa 594 and Star Red (Abberior) were used at 1/100 for STED imaging. Nuclei were counterstained with DAPI (Sigma-Aldrich). Images were obtained using a Laser Scanning confocal microscope (LSM780, Carl Zeiss) or through STED microscopy (TCS SP8 3X (Leica Microsystems), equipped with a pulsed white light laser to excite the Alexa594 at 561nm and the StarRed at 633nm. Both staining were depleted with the 775 nm pulsed laser (45% power for both, 4 and 6 accumulations respectively). All images were acquired at 700Hz through a Plan Apochromat 93x /1.3 NA Glycerol objective using the LAS X software (Leica Microsystems). All STED images had a 17-nm pixel size. All images were deconvoluted with Huygens Professional (version 18.10, Scientific Volume Imaging), using the CMLE algorithm (SNR :14, 40 iterations).

### Hypo-osmotic shock

Hypo-osmotic shock was performed by diluting growth medium with deionized water (1:9 dilution for 30 mOsm hypo-osmotic shock) for the indicated time.

### Shear stress

Cells were exposed to orbital shear stress with an orbital shaker (7 dynes/cm^2^) for the indicated time.

### Microtubule Purification

Microtubules were purified using the Taxol-Based Isolation of Tubulin from Cell as described^43^. Briefly, Adherent cells are washed and scraped in 1 ml of PBS. After centrifugation at 1200×g, cells are resuspended in MME buffer. The cell suspension is sonicated with a microtip probe seven times for 30 s with 30 s rest intervals on melted ice. The cell lysate is spun at 120,000×g (Beckman TL100 centrifuge) for 1h at 4°C. Cytosolic supernatants are incubated for 20 min at 37°C in the presence of 10 μM Taxol, 5% sucrose and 1 mM GTP. Samples are centrifuged at 80,000×g for 30 min at 37°C. Pellets are washed with 0.1 ml of warm MME buffer and resuspended in MME buffer containing 0.35 MNaCl and 10 μM Taxol on ice. After centrifugation at 80,000×g for 30 min at 37°C, microtubule pellets are frozen on dry ice and kept at −70°C until their use.

### Radioactivity

Posttranslational labeling was performed by incubating cells in normal growth medium containing 400 mCi/mL [^3^H]glutamine (PerkinElmer, 30-60 Ci/mmol) in the presence of (100 µg/ml) cycloheximide, a strong inhibitor of protein synthesis. These conditions resulted in the specific labeling of the polyglutamyl lateral chain (Eddé et al., 1992; Audebert et al., 1993). Purified microtubule from triplicate samples of Hela cells cultured under soft (1kPa) or stiff matrix (50kPa) for 48 hours were eluted by boiling in 2X sample buffer at 95°C for 10 min. The eluted fractions were analyzed by after SDS-PAGE. The radioactivity resulting from [3H]glutamine incorporation into tubulin was determined. Proteins were transferred onto nitrocellulose and the membrane was stained with Ponceau Red. Tubulin bands were then cut out and processed directly for liquid scintillation counting.

### Proteomic and stable carbon isotope proteomic analysis

Purified microtubule from triplicate samples of Hela cells cultured under soft (1kPa) or stiff matrix (50kPa) for 48 hours and treated with CB839 for 24 hours were analysed. To trace microtubule neo-glutamylation, cells were cultivated in DMEM containing 10% FBS, 4.5g.L^-1^ glucose, 2mM glutamine and then transferred into glutamine free (with 4.5g.L^-1^ glucose) DMEM containing 10% dialyzed FBS and supplemented with 2mM ^13^C_5_-glutamine for 2 hours in presence of (100 µg/ml) cycloheximide. Microtubules were purified according to the taxol-based purification procedure. Microtubule pellet were homogenized in 100 µl 50 mM ammonium bicarbonate and digested with thermolysin (Promega Corporation) at 65°c for 2h and at 37°c for an additional incubation time of 16h. Peptides were further dried under speed vacuum concentrator and resuspended in 20 µl Water/0.1% TFA (trifluoracetic Acid) for liquid chromatography (LC)–tandem mass spectrometry (MS/MS) analysis using an Orbitrap Fusion Lumos Tribrid mass spectrometer in-line with an Ultimate 3000 RSLCnano chromatography system (ThermoFisher Scientific). First, peptides (2µl, 10% of whole sample) were concentrated and purified on a pre-column from Dionex (C18 PepMap100, 2cm × 100µm I.D, 100Å pore size, 5µm particle size) in solvent A (0.1% formic acid in 2% acetonitrile). In the second step, peptides were separated on a reverse phase LC EASY-Spray C18 column from Dionex (PepMap RSLC C18, 50cm × 75µm I.D, 100Å pore size, 2µm particle size) at 300nL/min flow rate and 40°C. After column equilibration using 4% of solvent B (20% water - 80% acetonitrile - 0.1% formic acid), peptides were eluted from the analytical column by a two-step linear gradient (4-20% acetonitrile/H2O; 0.1% formic acid for 90min and 20-45% acetonitrile/H_2_O; 0.1% formic acid for 20min). For peptide ionization in the EASY-Spray nanosource, spray voltage was set at 2.2kV and the capillary temperature at 275°C. The Advanced Peak Determination (APD) algorithm was used for real time determination of charges states and monoisotopic peaks in complex MS spectra. The mass spectrometer was used in data dependent mode to switch consistently between MS and MS/MS. Time between Masters Scans was set to 3 seconds. MS spectra were acquired in the range m/z 400-1600 at a FWHM resolution of 120 000 measured at 200 m/z. AGC target was set at 4.0.10^5^ with a 50 ms maximum injection time. The most abundant precursor ions were selected and collision induced dissociation fragmentation at 35% was performed and analysed in the ion trap using the “Inject Ions for All Available Parallelizable time” option with a dynamic maximum injection time. Charge state screening was enabled to include precursors with 2 and 7 charge states. Dynamic exclusion was enabled with a repeat count of 1 and duration of 60s.

Raw files generated from mass spectrometry analysis were processed with Proteome Discoverer 1.4 (Thermo fisher Scientific). This software was used to search data via in-house Mascot server (version 2.4.1; Matrix Science Inc., London, UK) against the human tubulin proteins (Uniprot accession number : P68371, Q13509, Q13885, P04350, Q9BUF5, Q3ZCM7, Q9H853, Q9H4B7 for beta-tubulin and P0DPH7, Q9BQE3, Q6PEY2, P68366, Q71U36, Q9NY65, A6NHL2, P68363 for the alpha-tubulin). For alpha-tubulin subunit, sequences lacking the last carboxy-terminal tyrosine residues or the penultimate glutamate were also manually added to the fasta file to mimic putative posttranslational removal of the last or two last amino acids respectively^44,45.^ They were annoted P0DPH7_Y and P0DPH7_EY. To this fasta file, were added a subset of the swissprot human database corresponding to the 1159 first proteins among a total of 20,413 entries of the human SwissProt database (version 2019.11). Database search were done using the following settings: thermolysin cleavage before L, F, V, I, A or M amino acids with a maximum of 4 thermolysin miscleavage allowed, no static modifications. As glutamylation of tubulin is a complex modification with increasing length of glutamyl residues on the lateral chain of a glutamate residue of the main chain, we fixed parameters that will allow the identification of glutamyl chain bearing from 0 to 8 glutamyls units. As we were interested in the incorporation of neo-glutamylated residues, we looked for peptides that incorporated one heavy glutamyl ^13^Glu group ^13^C(5) H(7) N O(3)) in the modified chain. With these criteria, 8 dynamic modifications were used for database search:

|  |  |  |
| --- | --- | --- |
| $^{13}\text{Glu}$ | 134.059367 | $^{13}\text{C}(5)$ H(7) N O(3) |
| $\text{Glu}^{13}\text{Glu}$ | 263.101960 | $^{13}\text{C}(5)$ C(5) H(14) N(2) O(6) |
| $\text{GluGlu}^{13}\text{Glu}$ | 392.144554 | $^{13}\text{C}(5)$ C(10) H(21) N(3) O(9) |
| $\text{GluGluGlu}^{13}\text{Glu}$ | 521.187147 | $^{13}\text{C}(5)$ C(15) H(28) N(4) O(12) |
| $\text{GluGluGluGlu}^{13}\text{Glu}$ | 666.224654 | $^{13}\text{C}(5)$ C(20) H(35) N(5) O(16) |
| $\text{GluGluGluGluGlu}^{13}\text{Glu}$ | 779.272333 | $^{13}\text{C}(5)$ C(25) H(42) N(6) O(18) |
| $\text{GluGluGluGluGluGlu}^{13}\text{Glu}$ | 908.314926 | $^{13}\text{C}(5)$ C(30) H(49) N(7) O(21) |
| $\text{GluGluGluGluGluGluGlu}^{13}\text{Glu}$ | 1037.357519 | $^{13}\text{C}(5)$ C(35) H(56) N(8) O(24) |

A peptide mass tolerance of 6 ppm and a fragment mass tolerance of 0.2 Da were allowed for search analysis. Only peptides with higher Mascot threshold (identity) were selected. False discovery rate was set to 1% for protein identification. For all experiments, the quantity of the glutamatylated C-terminal tubulins tail identified was normalized by the quantity of corresponding C-terminal tubulin tail identified. Number of PSM (Peptide Spectrum Matches) values from Proteome Discoverer results were used for spectral counting quantification. Peptides VDSVEGEGEEEGEEY, LEKDYEEVGVDSVEAE from alpha 1 and alpha 3d carboxy terminal tubulin sequences and peptides LVSEYQQYQDATAEEEEDEEYAEEE, from beta-8 tubulin isotype showed ^13^Glu neo incorporation.

### Western blot assays

Forty-eight hours after plaiting, cells were lysed in RIPA buffer (Pierce) or directly in Laemmli’s buffer. After denaturation, protein lysates were resolved by SDS-PAGE and transferred onto a PVDF membrane (Millipore). Membranes were blocked with 2% BSA in TBS tween20 0.1% and incubated in the presence of the primary and then secondary antibodies. After washing, immunoreactive bands were visualized with ECL (Millipore) and analyzed on Fusion-FX Imager (Vilber). ImageJ software was used to quantify band intensity and the ratios of proteins of interest were normalized to Hsp90 (loading control). For the analysis, Glu-Tubulin levels were quantified by calculating the ratio between Glu-Tubulin and Tubulin, both normalized to Hsp90 signal as follows: (Glu-Tubulin /Hsp90^Glu-Tubulin^) / (Tubulin /Hsp90 ^Tubulin^). Mean expression in controls was assigned a fold change of 1, to which relevant samples were compared.

### RNA isolation, RT-PCR and q-PCR

Total RNA was extracted by TRIzol reagent according to the manufacturer’s instructions (Invitrogen). RNA quantity and quality were determined using NanoDrop™One Spectrophotometer (Thermo Scientific). One microgram of total RNA was reverse transcribed to generate cDNA (A3500, Promega). cDNA was amplified via fluorescently labeled Taqman primer sets using an Applied Biosystems 7900HT Fast Real Time PCR device. Fold change of RNA species was calculated using the formula (2^-^ΔΔ^Ct^), normalized to RPLP0 expression. All real-time RT-PCR assays were performed in triplicate with three independent experiments. Primers are provided in Table:

|  | Taqman Catalog Number |
| --- | --- |
| RPLP0 | Hs99999902 |
| TTLL1 | Hs00923132 |
| TTLL4 | Hs01565335 |
| TTLL5 | Hs00209404 |
| TTLL6 | Hs00543401 |
| TTLL7 | Hs01114458 |
| TTLL9 | Hs01082287 |
| TTLL11 | Hs00879398 |
| TTLL13 | Hs01050281 |
| CCP1 | Hs00999967 |
| CCP4 | Hs00328701 |
| CCP2 | Hs01050124 |
| CCP3 | Hs01570549 |
| CCP6 | Hs00901829 |
| CCP5 | Hs00222447 |

### Atomic force microscopy

Cells were first washed twice with 3 ml of Liebovitz’s medium (Life Technologies) supplemented with heat-inactivated 10% Fetal Bovine Serum (FBS), then covered with 3 ml of the same medium. The mechanical properties of samples were studied using a BioScope Catalyst atomic force microscope (Bruker Nano Surfaces, Santa Barbara, CA, USA) equipped with a Nanoscope V controller and coupled with an optical microscope (Leica DMI6000B, Leica Microsystems Ltd., UK). For each sample, at least 30 cells were analyzed using the “Point and Shoot” method, collecting at least 150 force-distance curves at just as many discrete points (on average 5 points for each cell in the perinuclear area). The experiments were performed using a probe with a borosilicate glass spherical tip (5 μm diameter) and a cantilever with a nominal spring constant of 0.06 N/m (Novascan Technologies, Ames, IA USA). After determining both the deflection sensitivity of the system in the Leibovitz’s medium using a clean Willco Glass Bottom Dish and the spring constant of the cantilever by the thermal tune method, force-distance curves were collected on samples using a velocity of 2 μm/s, in relative trigger mode and by setting the trigger threshold to 1 nN. The apparent Young’s modulus was calculated using the NanoScope Analysis 1.80 software (Bruker Nano Surfaces, Santa Barbara, CA, USA) applying to the force curves, after the baseline correction, the Hertz spherical indentation model using a Poisson’s ratio of 0.5. Only the force curves having their maximum value at 1 nN were considered for the analysis. For each force curve, the force fit boundaries to perform the fit were chosen between 50 and 250 pN and only the apparent Young’s modulus values corresponding to a fit with R^2^> 0.80 were accepted.

### Traction force microscopy

Contractile forces exerted by cells on different stiffness gels were assessed by traction force microscopy essentially as describe^5^. Briefly, polyacrylamide substrates with shear moduli of 1kPa or 12 kPa conjugated with fluorescent bead latex microspheres (0.5 μm, 505/515 nm ex/em) were purchased from Matrigen. After transfection, cells were plated on fluorescent bead–conjugated discrete stiffness gels and grown for 24 hours. Images of gel surface– conjugated fluorescent beads were acquired for each cell before and after cell removal using an Axiovert 200M motorized microscope stand (Zeiss) and a ×32 magnification objective. Tractions exerted by cells were estimated by measuring bead displacement fields, computing corresponding traction fields using Fourier transform traction microscopy, and calculating root-mean-square traction using the PIV (Particle Image Velocimetry) and TFM (Traction force microscopy) package on ImageJ^46^. To measure baseline noise, the same procedure was performed on a cell-free region.

### FRAP

Celllight GFP-Tubulin (Thermo, C10613) were add to cells sixteen hours before experiment or Tubulin Tracker Green were add to cells thirty minutes before experiment to labelled polymerized tubulin in live cells. FRAP experiment were performed on Laser Scanning Confocal Microscope (LSM780 Carl Zeiss) equipped with a heated stage maintained at 37°C and the whole power of its 30mW 488 nm Argon laser. Fluorescence intensity variations in microtubule were analyzed using the FRAP module of ZEN software (Carl Zeiss). Mobile fraction (in %) and half-time of recovery (in seconds) values were extracted for each experiment.

### Circularity measurement

The circularity index of cell shape was quantified as previously described^47^. For application of circularity measurements of cells, a freehand selection option in ImageJ software was used to delineate cell outline. Circularity index values were assigned to cell outlines by the ImageJ circularity plugin where circularity = 4p(area/perimeter2).

### Cell adhesion

Cells were detached, washed three times with PBS and 3×10^4^ cells were seeded per well of 24 well-plate previously coated with collagen. After 15 min of adhesion, cells were washed three times with PBS, fixed with PFA 4% and stained with DAPI. Cell adhesion were determined by counting all cells on the total area with Cytation5 Biotek System. Nuclei were counted and analyzed using Image J software.

### Cell proliferation

Cells were detached, washed three times with PBS and 3×10^4^ cells were seeded per well of 24 well-plate previously coated with collagen. After 24 hours of proliferation, cells were washed three times with PBS, fixed with PFA 4% and stained with dapi. Cell proliferation was determined by imaging all the cells on the total area of the wells with a Image plate-reader (Cytation5, Biotek). Nuclei were counted and analyzed using ImageJ software.

### Microtubules alignment analysis

Cell alignment was calculated from microscopy images based on Local Gradients orientation method analysis using the directionality plug-in available through ImageJ^48^. A histogram with the peak of the dominant orientation was produced for each representative image. A completely flat histogram represents an isotropic orientation^49^

### Live imaging and cell velocity/speed measurement

Forty-two hours after siRNA transfection, cells were collected, reseeded on 24 wells plate, allowed to adhere, and migration was monitored at the earliest 4 h later. Cell migration was monitored by brightfield phase contrast microscopy using an Axiovert 200M videomicroscope (Carl Zeiss) with a 10X/0.3 Ph1 objective every 10 min for 2 h in a 37°C temperature-controlled environmental chamber. Persistence and velocity were analyzed using the Manual Tracking plugin of Fiji^48^.

### Glutamylated-tubulin intensity

The mean intensity of Glu-Tubulin (Alexa488 fluorescence) in the cell area was measured with ImageJ software (NIH) as above on z-projections, after background correction.

### EB1 Comet Imaging Assays and Analysis

The cells were imaged 24h post transfection to measure the dynamicity of microtubules ends labeled with GFP-EB1. Images were acquired with a 488nm excitation on a spinning disk confocal microscope (Ultraview Vox, Perkin Elmer) equipped with a TiE Eclipse (Nikon Instruments) stand through a CFI PlanApochromat 100X/1.4 NA oil objective every 0.5 s during 1 min for each time-lapse. All the image analysis of the sequences was performed using MTrack plugin of Fiji^50^

### Medium metabolite measurements

For kinetics of metabolite secretion by HeLa cells, triplicate samples of subconfluent HeLa cultured under soft (1kPa) or stiff (12kPa and 50kPa) condition were changed to fresh DMEM with 10% FBS, which was allowed to condition for 48 h. Metabolites were then extracted from conditioned medium by adding ice cold 100% MeOH to a final concentration of 80% MeOH. Medium collected from cell-free plates after 48 h incubation was used as the baseline control to calculate the consumption or production of each metabolite, which was further normalized by the proliferation rate. The cell numbers were measured from duplicate treatment plates to determine the proliferation rate, and the metabolite flux was determined with the following formula:

Uptake/secretion rate = Δ metabolite / (Δ time * average cell number)

Average cell number = Δ cell number / (growth rate* Δ time)

Uptake/secretion rate = (Δ metabolite / Δ time) * (growth rate * Δ time / Δ cell number) = (Δ metabolite / Δ cell number) * growt rate

Growth rate [1/h] = LN (cell number T1) – LN (cell number T0) / time (T1)-time (T0)

### Targeted LC-MS

Metabolite extraction was performed essentially as described with minor modifications^5,51^. Briefly, metabolites were extracted from cultured cells on dry ice using 80% aqueous methanol precooled at –80°C. Supernatants were extracted with 4 volumes of 100% methanol precooled at –80°C for 4 hours at –80°C. An internal standard, [^13^C_4_]-2-oxoglutarate ([^13^C_4_]-2OG) (Cambridge Isotope Laboratories), was added during metabolite extraction. Insoluble material from both cell and supernatant extractions was removed by centrifugation at 20,000 g for 15 minutes at 4°C. The supernatant was evaporated to dryness by SpeedVac at 42 °C, the pellet was resuspended in LC-MS water, and metabolites were analyzed by LC-MS.

LC-MS analysis was performed on a Vanquish ultra-high performance liquid chromatography system coupled to a Q Exactive mass spectrometer (Thermo) that was equipped with an Ion Max source and HESI II probe. External mass calibration was performed every seven days. Metabolites were separated using a ZIC-pHILIC stationary phase (150 mm × 2.1 mm × 3.5 mm; Merck) with guard column. Mobile phase A was 20 mM ammonium carbonate and 0.1% ammonium hydroxide. Mobile phase B was acetonitrile. The injection volume was 1 μL, the mobile phase flow rate was 100 μL/min, the column compartment temperature was set at 25 °C, and the autosampler compartment was set at 4 °C. The mobile phase gradient (%B) was 0 min, 80%; 5 min 80%; 30 min, 20%; 31 min, 80%; 42 min, 80%. The column effluent was introduced to the mass spectrometer with the following ionization source settings: sheath gas 40, auxillary gas 15, sweep gas 1, spray voltage +/- 3.0 kV, capillary temperature 275 °C, S-lens RF level 40, probe temperature 350 °C. The mass spectrometer was operated in polarity switching full scan mode from 70-1000 m/z. Resolution was set to 70,000 and the AGC target was 1×10^6^ ions. Data were acquired and analysed using TraceFinder software (Thermo) with peak identifications based on an in-house library of authentic metabolite standards previously analysed utilizing this method. For all metabolomic experiments, the quantity of the metabolite fraction analysed was adjusted to the corresponding cell number calculated upon processing a parallel experiment.

### Measurements of metabolite levels by kits

The levels of selected metabolites were measured by commercial kits to confirm the results of metabolic profiling. These include the glutamate colorimetric assay kit (BioVision) and the glutamine colorimetric assay kit (BioVision). The manufacturers’ protocols were followed. Cell number was determined in concurrent experiment run in parallel, averaged per condition, and the metabolite consumption/production rates were calculated per cell.

### Glutaminase activity assay

According to the manufacturer instructions (Glutaminase Microplate Assay Kit, Cohesion Biosciences), flash frozen tissue (0.1g/sample) or cells (1×10^6^ cells) was homogenized in 1mL of assay buffer on ice and centrifuged at 8000g 4°C for 10 min. Protein concentration was determined by Bradford assay. Samples, normalized to total protein (100μg) or cell numbers, were incubated with kit reagents for 1 hr at 37°C, and absorbances were measured at 420nm.

### Statistical analysis

All analyses were performed using Prism 6.0 software (GraphPad Inc.). A two-tailed t-test was used if comparing only two conditions. For comparing more than two conditions, one-way ANOVA was used with: Bonferroni’s multiple comparison test or Dunnett’s multiple comparison test (if comparing all conditions to the control condition). Significance of mean comparison is marked on the graphs by asterisks. Error bars denote SEM.

### Data availability

All the data are available upon reasonable request.

## Acknowledgements

We thank the B. Mari team members for advice and discussions as well as F. Aguila for artwork. The author acknowledge the “Microscopie Imagerie Côte d’Azur” (MICA), GIS-IBISA multi-sites platform and particularly it’s IPMC, C3M and IRCAN (Molecular and Cellular Imaging facility PICMI) partners. This platform is supported by the GIS IBiSA, Conseil Départemental 06, Région PACA ARC, Cancerôpole PACA,)”. Proteomic analyses were performed at the mass spectrometry facility of Marseille Proteomics supported by IBISA, Plateforme Technologique Aix-Marseille, Canceropôle PACA, Région Sud Provence-Alpes-Côte d’Azur, Fonds Européen de Développement Régional (FEDER) and Plan Cancer.

## Author contributions

ST and TB conceived and designed the experiments. ST, SA, IB, WMO and TB performed the majority of the experiments. WMO, BM, and TB, provided the experimental infrastructure. CL generated tubulin mutants. FB performed and analysed FLIM experiments. SA performed STED experiments. SP performed and analysed AFM experiments. S. Audebert performed and analysed proteomic experiments. ST, WMO, BM, SYC, and TB wrote the manuscript. All authors participated in interpreting the results and revising the manuscript.

## Extended data for

**Extended figure 1:**
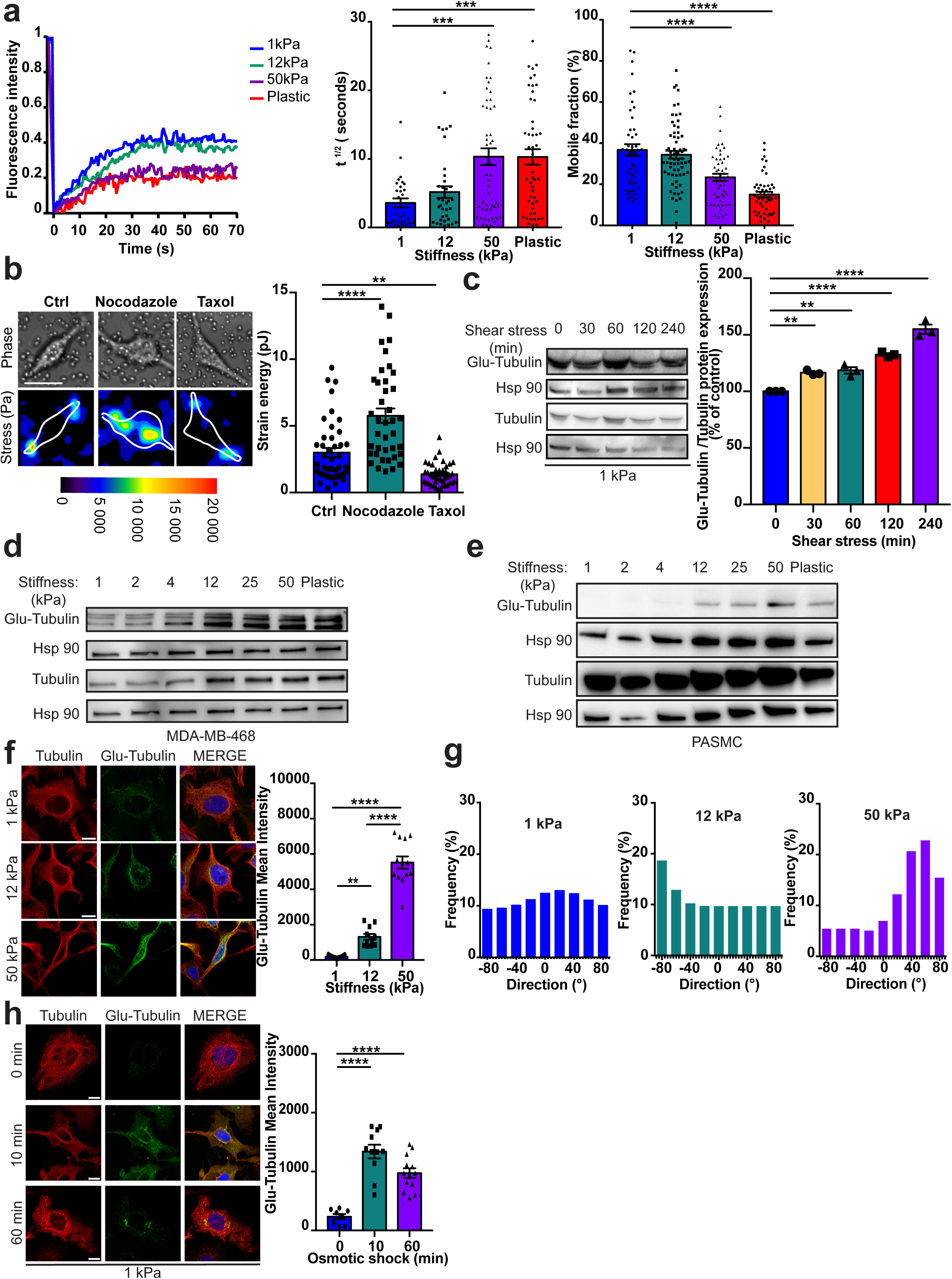
Mechanical cues increase microtubules glutamylation and alter microtubules dynamics to force cell mechanics. **(a)** Representative FRAP curves (left) and quantification of diffusion rate (t ½) and mobile fraction (right) of endogenous Tubulin labeled with Oregon Green™ 488 Taxol, Bis-Acetate in HeLa cells plated on different stiffness hydrogel (1, 12, 50 kPa) or plastic (n>45 cells). **(b)** Representative heat map (left) and quantification (right; n>15 cells from n=3 independent experiments) showing contractile forces generate by cells plated on 12kPa hydrogel and treated with Nocodazole or Taxol. **(c)** Immunoblot and quantification (n=3 independent experiments) of Glu-Tubulin in HeLa cells plated on 1 kPa hydrogel and after shear stress for the indicated times. Hsp90 was used as a loading control. **(d**,**e)** Immunoblot of Glu-Tubulin in MDA-MB-468 cells (d) and primary pulmonary arterial smooth muscle cells (PASMCs; e) plated on the indicated substrate. **(f**,**h)** Representative confocal images of Tubulin and Glu-Tubulin localization in HeLa cells plated on different stiffness hydrogel (1, 12, 50 kPa) or plated on 1kPa hydrogel and after osmotic stress for the indicated times. Nuclei were stained with DAPI (Blue) on the MERGE image. Quantification (right) of Glu-Tubulin intensity in the different condition. At least 50 cells per condition. Scale bar=10 µm. **(g)** Representative alignment of microtubule in HeLa cells plated on different stiffness hydrogel (1, 12, 50 kPa). At least 10 cells per condition. n=3 independent experiments; **P<0.01; ***P<0.001; ****P<0.0001; **(a-c)** Bonferroni’s multiple comparison test; data are mean ± s.e.m.

**Extended figure 2:**
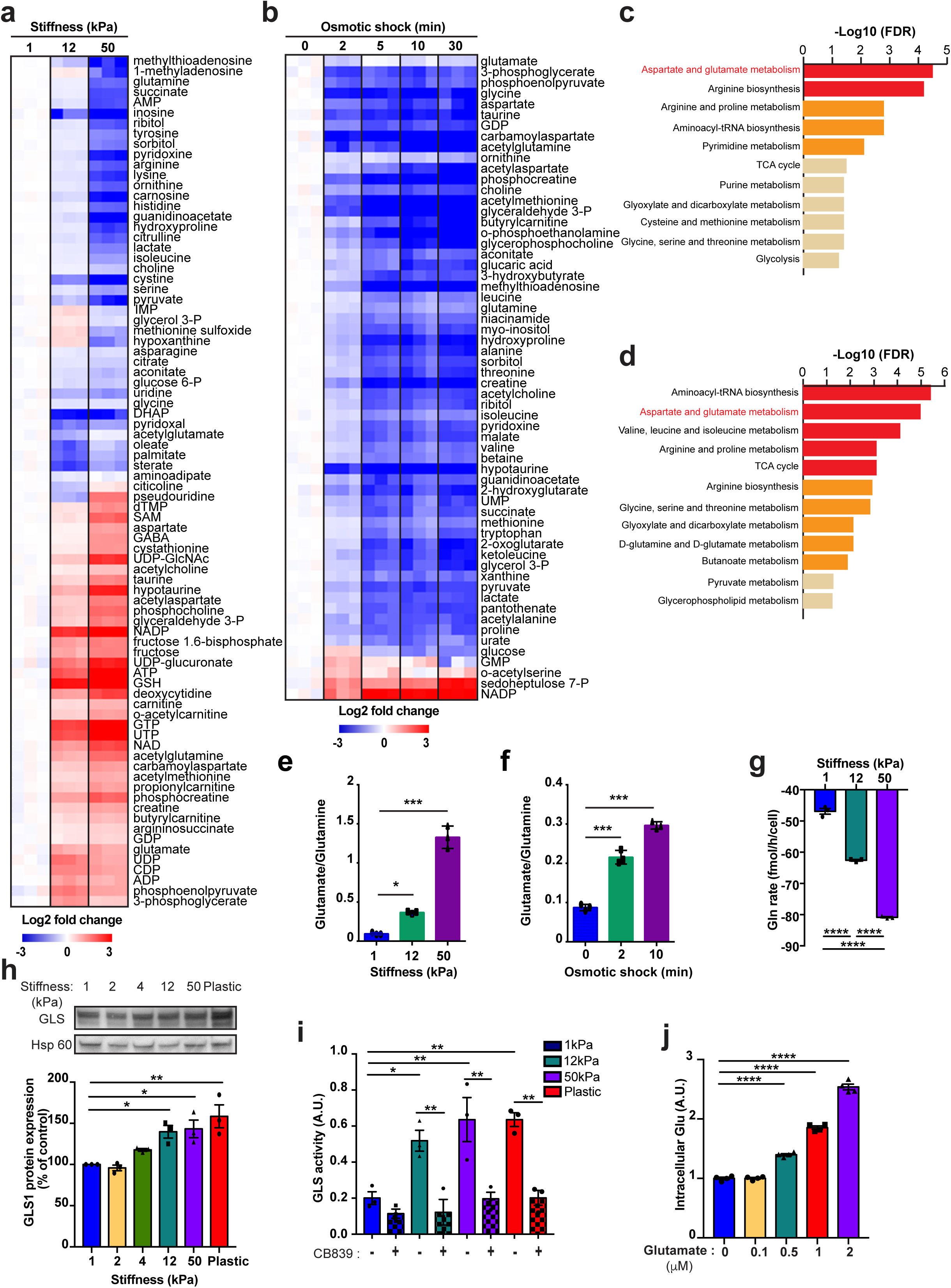
Mechanoactivation of GLS-dependent glutamine catabolism sustain the intracellular glutamate pool. **(a-d)** Heatmap **(a**,**b)** and pathway enrichment analysis **(c**,**d)** of significantly (FDR<1%; P<0.05) modulated intracellular metabolites in cells plated on the indicated substrate **(a**,**c)** or after hypo-osmotic shock **(b**,**d).** Red: metabolite from aspartate glutamate pathway. **(e**,**f)** Glutamate/Glutamine ratio in cells plated on the indicated substrate **(e)** or after hypo-osmotic shock **(f)**. Data are extract from (a-b) analysis. **(g)** Glutamine flux of HeLa cells plated on hydrogel of the indicated stiffness (n=3). **(h-i)** HeLa cells plated on different stiffness hydrogel (1, 12, 50 kPa) or plastic. **(h)** Immunoblot and quantification (n=3) of GLS. **(i)** Measurement of GLS activity in cells treated with CB839 for 24h. **(j)** Intracellular glutamate level of HeLa cells overexpressing SLC1A3 cultivated on 1kPa hydrogel and in presence of various glutamate concentration (n=3).

**Extended figure 3:**
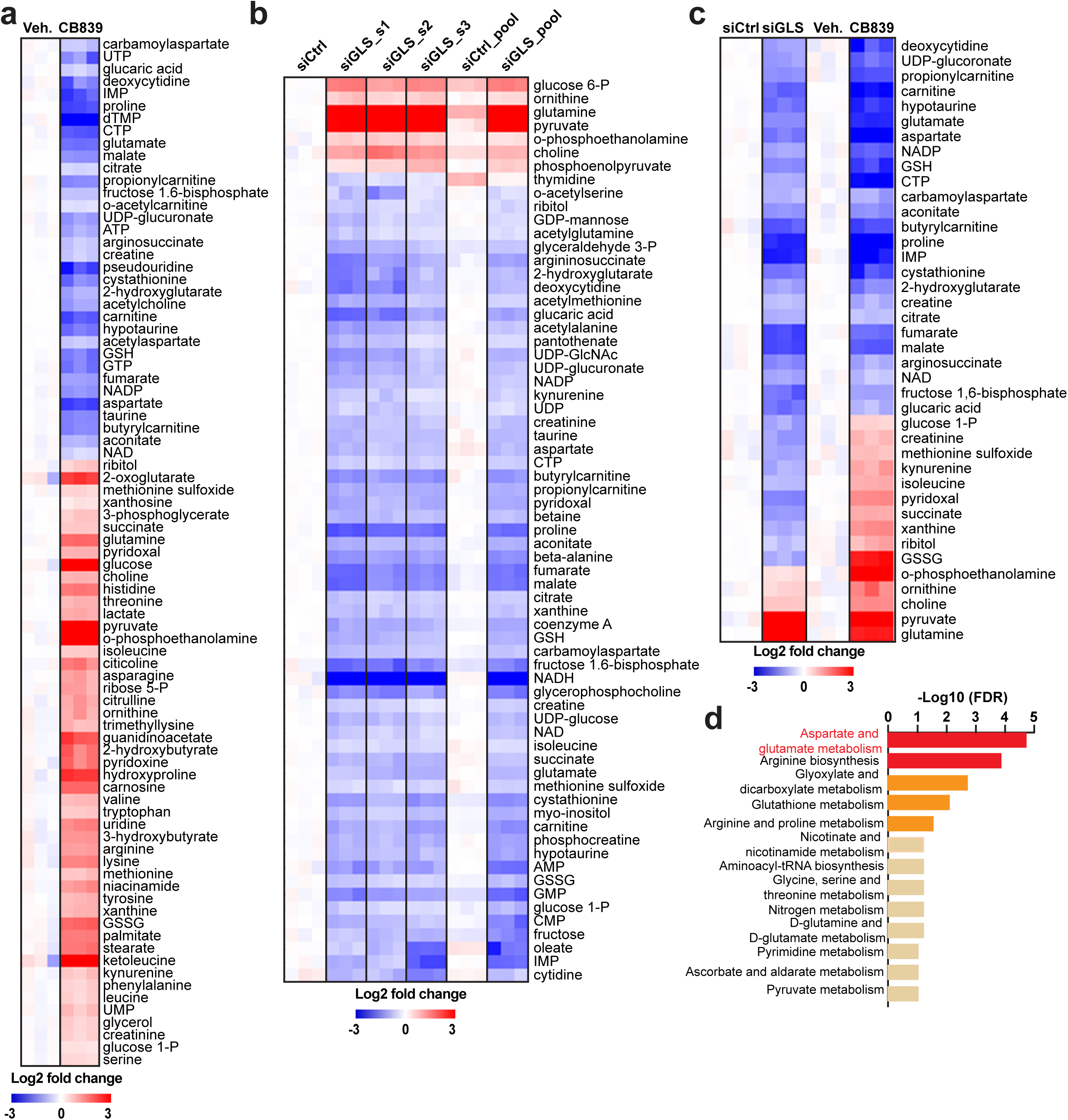
Either genetic or pharmacologic inhibition of GLS rewired glutamate metabolism. **(a-d)** Heatmap **(a**,**c)** and pathway enrichment analysis **(d)** of significantly (FDR<1%; P<0.05) modulated intracellular metabolites in cells following indicated treatments. Red: metabolite from aspartate glutamate pathway.

**Extended figure 4:**
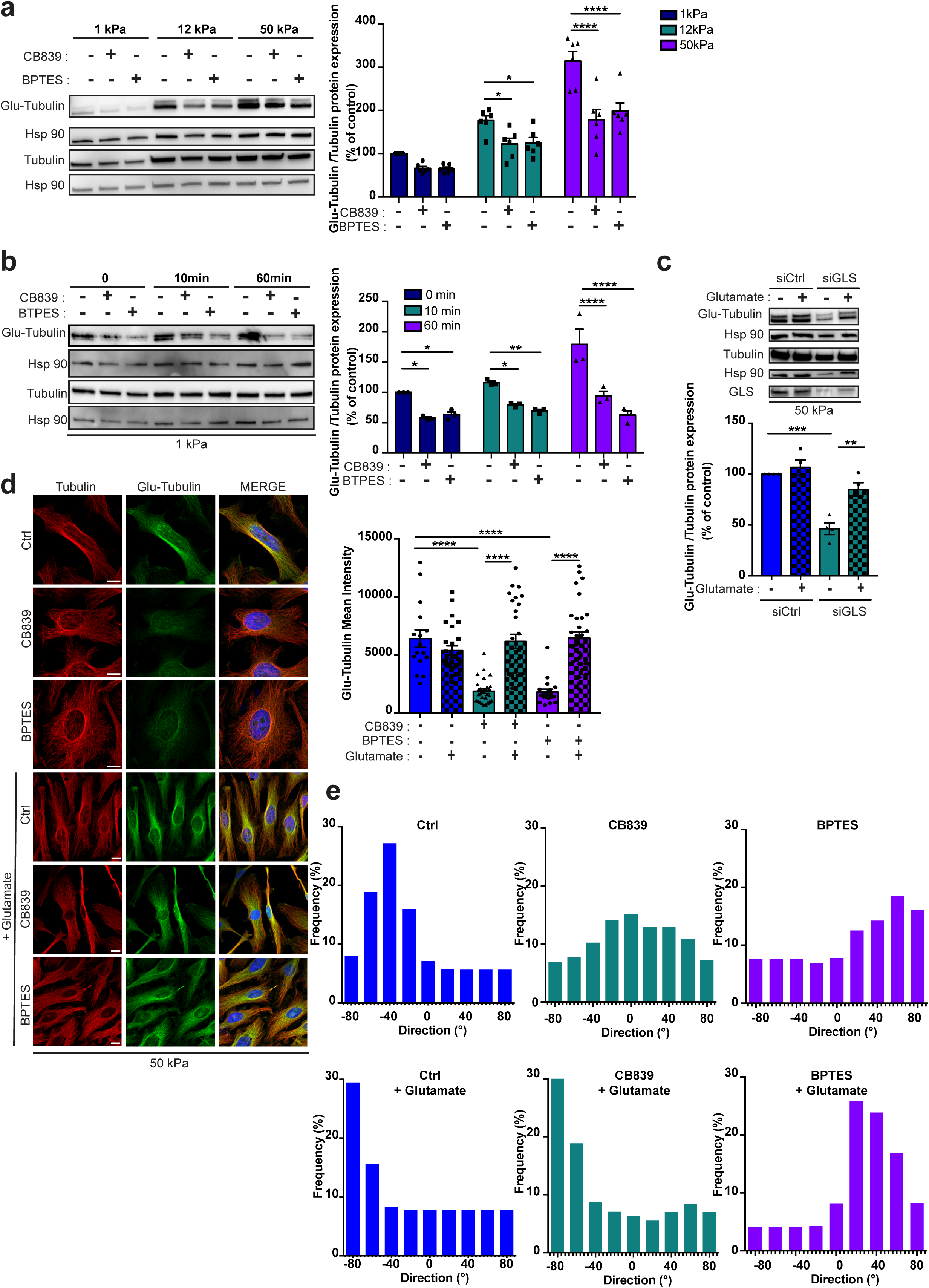
Mechanoactivation of GLS-dependent glutamine catabolism sustain microtubules glutamylation under mechanical stresses. **(a-b, d-e)** HeLa cells plated on the indicated substrate and treated with CB839 or BPTES. **(a)** Immunoblot and quantification (n=3 independent experiments) of Glu-Tubulin in cells. **(b)** Immunoblot and quantification (n=3) of Glu-Tubulin in HeLa cells after osmotic stress for the indicated times. Hsp90 was used as a loading control. **(c)** Immunoblot and quantification (n=3) of Glu-Tubulin in HeLa cells transfected with the siRNA GLS in presence of glutamate for 24h. **(d)** Representative confocal images (left) and quantification (right; n>50 cells from n=3 independent experiments) of Glu-Tubulin and Tubulin in presence of glutamate for 24h. Nuclei were stained with DAPI (Blue) on the MERGE image.Scale bar=10 µm. At least 50 cells per condition from 3 independent experiments. Scale bar=10 µm. **(e)** Representative alignment of microtubule in cells in presence of glutamate for 24h. At least 10 cells per condition from n=3 independent experiments; *P<0.05; **P<0.01; ***P<0.001; ****P<0.0001; **(a**,**b)** two-way ANOVA and Dunnett’s multiple comparisons test; **(c**,**d)** Bonferroni’s multiple comparison test;; data are mean ± s.e.m.

**Extended figure 5:**
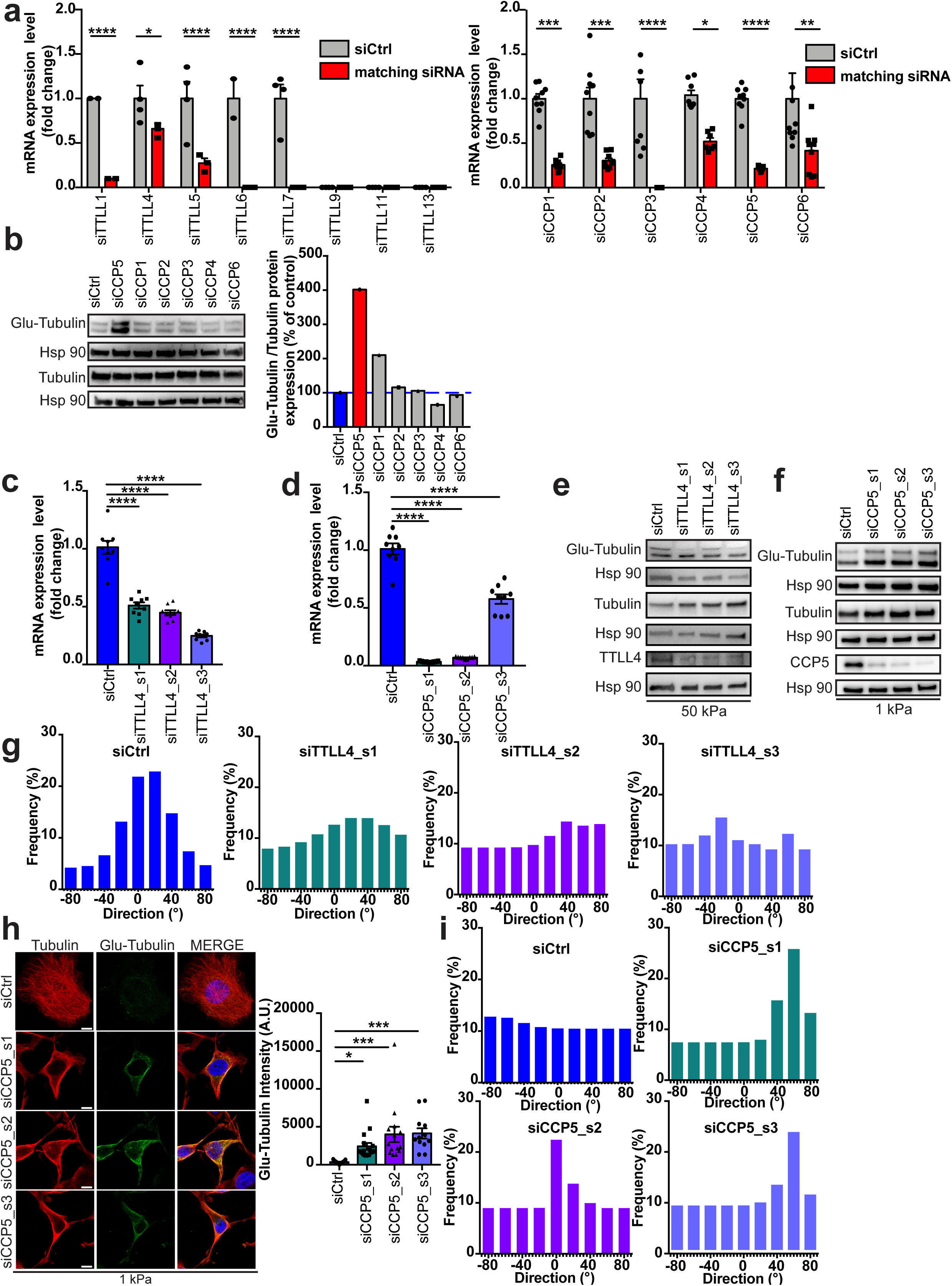
Balanced microtubules glutamylation by TTL4 and CCP5 organize the microtubule lattice. **(a-i)** HeLa cells were transfected with the indicated siRNA (control, siCtrl; TTLL and CCP, siRNA smarpool or siRNA single, _s1, s2, s3) for 48h. **(a**,**c-d)** As demonstrated by RT-qPCR, effective siRNA knockdown was achieved in Hela cells. For each gene transcript, mean expression in control groups (siCtrl) were assigned a fold change of 1, to which relevant samples (transfected with a siRNA specific to that gene) were compared**. (b**,**e-f)** Immunoblot **(b**,**e-f)** and quantification **(b)** of Glu-Tubulin in cells. **(g, i)** Representative alignment of microtubule in cells in plated on 50 kPa **(g)** or 1kPa **(i)** hydrogels. **(h)** Representative confocal images (left) and quantification (right; n>50 cells from 3 independent experiments) of Glu-Tubulin and Tubulin in cells plated on 1kPa hydrogel. Scale bar=10 µm. At least 10 cells per condition from n=3 independent experiments; *P<0.05; **P<0.01; ***P<0.001; ****P<0.0001; **(a)** two-way ANOVA and Dunnett’s multiple comparisons test; **(c-d, h)** Bonferroni’s multiple comparison test; data are mean ± s.e.m.

**Extended figure 6:**
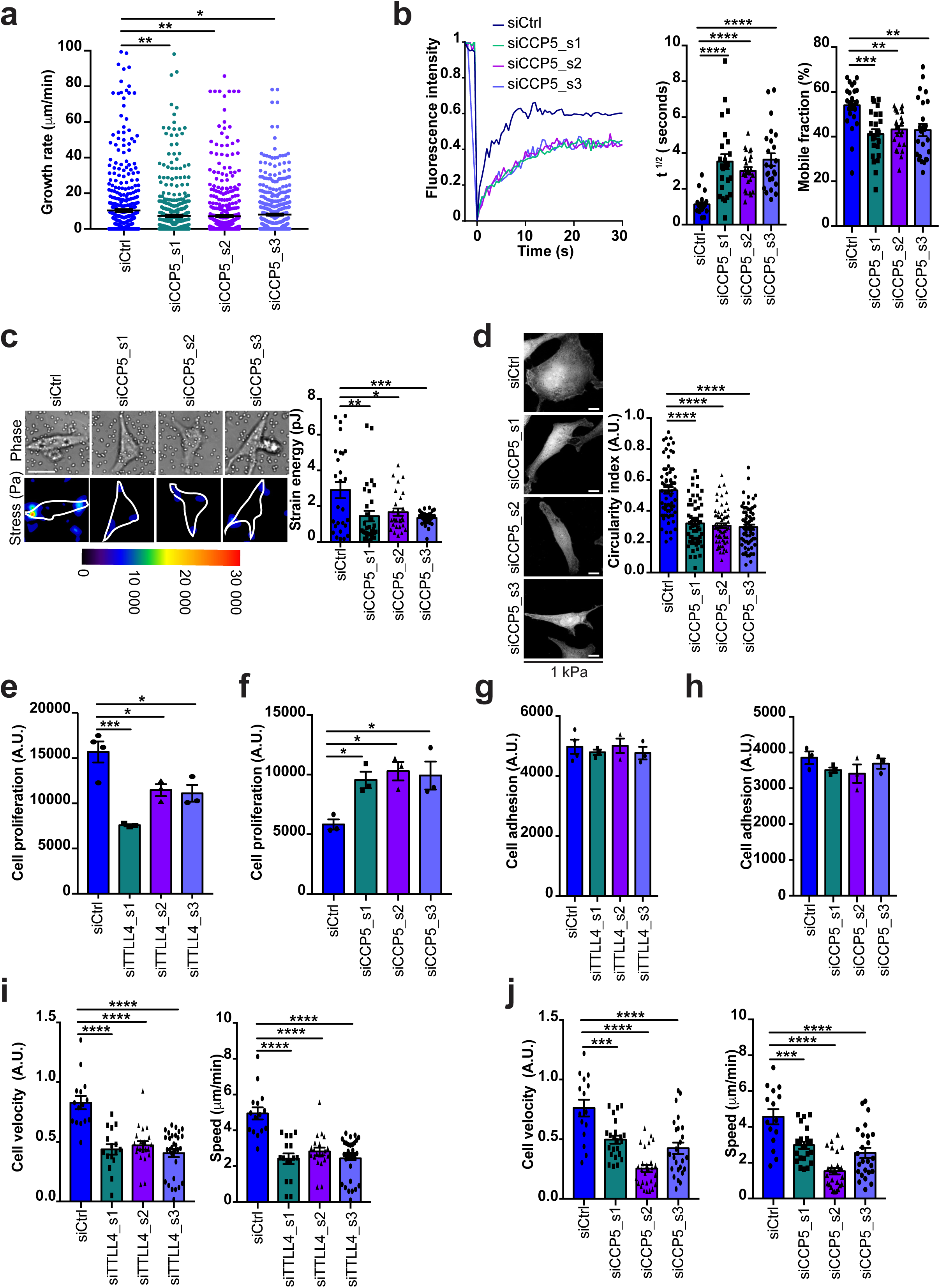
Microtubules glutamylation is orchestrated by TTLL4 and CCP5 to adjust cell mechanics and sustain cell mechanic-dependent activities. **(a-j)** HeLa cells were transfected with the indicated siRNA (control, siCtrl; TTLL and CCP, siRNA single, _s1, s2, s3) for 48h. **(a)** Representative kymographs (left) and growth rates quantification (right) of EB1-GFP in cells plated on 1kPa hydrogel. n>500 comets. Scale bar=1 µm. **(b)** Representative FRAP curves (left) and quantification of diffusion rate (t ½) and mobile fraction (right) of GFP-Tubulin in cells plated on 1kPa hydrogel (n>45 cells). **(c)** Representative heat map (left) and quantification (right) showing contractile forces generate by cells plated on 12kPa hydrogel. **(d)** Representative confocal images (left) and quantification (right) of circularity index of cells plated on 1kPa hydrogel. Scale bar=10 µm. **(e-f)** Proliferation rate of cells plated on 50kPa **(e)** or 1kPa **(f)** hydrogel. **(g-h)** Measurement of cell adhesion on cells plated on 50kPa **(g)** or 1kPa **(h)** hydrogel. **(i-j)** Cell velocity (left) and speed (right) of cells plated on 50kPa **(i)** or 1kPa **(j)** hydrogel. In all the panels n>50 cells from 3 independent experiments were analyzed. *P<0.05; **P<0.01; ***P<0.001; ****P<0.0001; **(a-j)** Bonferroni’s multiple comparison test; data are mean ± s.e.m.

**Extended figure 7:**
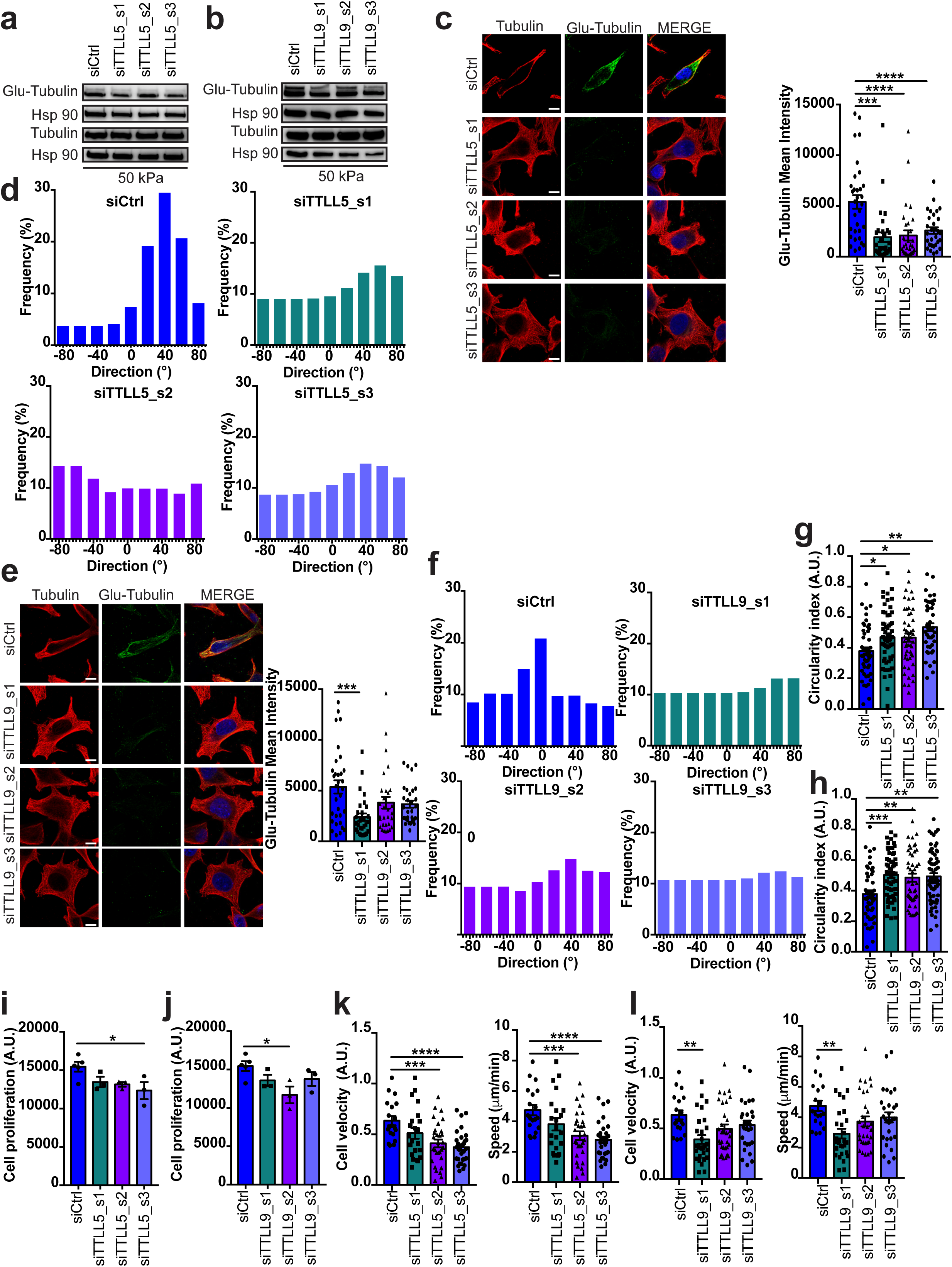
TTLL5 orTTLL9 cell depletion decreased MT glutamylation, reorganize the microtubule lattice and affect cell mechanic-dependent cell functions. **(a-l)** HeLa cells were transfected with the indicated siRNA (control, siCtrl; TTLL and CCP, siRNA single, _s1, s2, s3) for 48h and plated on 50kPa hydrogel. **(a**,**b)** Immunoblot of Glu-Tubulin in cells. **(c**,**e)** Representative confocal images (left) and quantification (right; n>50 cells from 3 independent experiments) of Glu-Tubulin and Tubulin in cells. Scale bar=10 µm. At least 10 cells per condition from n=3 independent experiments. **(d, f)** Representative alignment of microtubule in cells. **(g**,**h)** Quantification of circularity index of cells. **(i-j)** Proliferation rate of cells. **(k-l)** Cell velocity (left) and speed (right) of cells. In all the panels n>50 cells from 3 independent experiments were analyzed. *P<0.05; **P<0.01; ***P<0.001; ****P<0.0001; **(c**,**e**,**g-l)** Bonferroni’s multiple comparison test; data are mean ± s.e.m.

**Extended figure 8:**
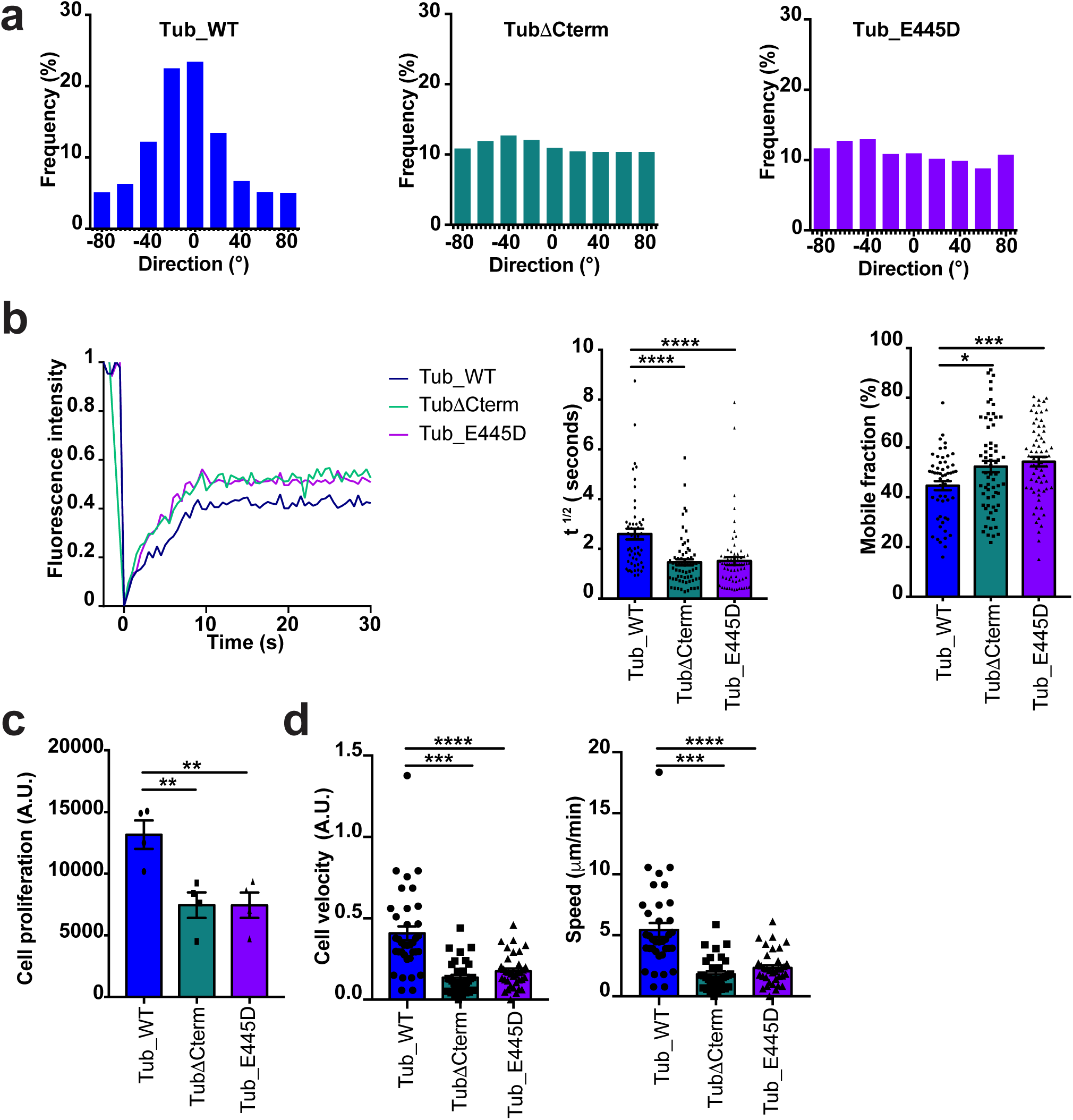
C-terminal tubulin tail mutants increase microtubule dynamics and modulate cell mechanic-dependent activities. **(a-d)** HeLa cells were transfected with TUBA1A constructs and plated on 50kPa hydrogel. **(a)** Representative alignment of microtubule in cells. **(b)** Representative FRAP curves (left) and quantification of diffusion rate (t ½) and mobile fraction (right) of GFP-Tubulin in cells. **(c)** Proliferation rate of cells. **(d)** Cell velocity (left) and speed (right) of cells. In all the panels n>50 cells from 3 independent experiments were analyzed. *P<0.05; **P<0.01; ***P<0.001; ****P<0.0001; **(c-d)** Bonferroni’s multiple comparison test; data are mean ± s.e.m.

